# SARS-CoV-2 Spike evolution influences GBP and IFITM sensitivity

**DOI:** 10.1101/2022.03.07.481785

**Authors:** Dejan Mesner, Ann-Kathrin Reuschl, Matthew V.X Whelan, Taylor Bronzovich, Tafhima Haider, Lucy G. Thorne, Greg J. Towers, Clare Jolly

**Author notes:** Corresponding author: Clare Jolly. These authors contributed equally (Dejan Mesner, Ann-Kathrin Reuschl).

## Abstract

SARS-CoV-2 spike requires proteolytic processing for viral entry. The presence of a polybasic furin-cleavage site (FCS) in spike, and evolution towards an optimised FCS by dominant variants of concern (VOCs), are linked to enhanced infectivity and transmission. Here we show that interferon-inducible antiviral restriction factors Guanylate binding proteins (GBP) 2 and 5 interfere with furin-mediated cleavage of SARS-CoV-2 spike and inhibit the infectivity of early-lineage Wuhan-Hu-1, while VOCs Alpha and Delta have evolved to escape restriction. Strikingly, we find Omicron is unique amongst VOCs, being restricted by GBP2/5, and also IFITM1, 2 and 3. Replacing the spike S2 domain in Omicron with Delta shows S2 is the determinant of entry route and IFITM sensitivity. We conclude that VOC evolution under different selective pressures has influenced sensitivity to spike-targeting restriction factors, with Omicron selecting spike changes that not only mediate antibody escape, and altered tropism, but also sensitivity to innate immunity.

## Introduction

SARS-CoV-2 infects cells by binding of the viral spike (S) protein to the angiotensin converting enzyme 2 (ACE2) receptor on host cells ^1^. For fusion to proceed after ACE2 binding, spike must be cleaved by host cell proteases to become activated and fusion-competent. In this step-wise process spike is pre-processed by furin-like proteases in virus-producing cells at the S1/S2 junction ^2, 3^, followed by a second cleavage event at the S2’ site mediated by TMPRSS2 protease at the target cell surface ^1^, releasing the fusion peptide and allowing for viral fusion at the plasma membrane. Alternatively, TMPRSS2-independent endocytic uptake can occur in some cell types resulting in spike being cleaved and activated for fusion by endosomal cathepsin proteases ^2^. The polybasic furin cleavage site (FCS) Arg-Arg-Ala-Arg (or RRAR) motif that is targeted by furin is absent in closely related coronaviruses, including the closest relatives of SARS-CoV-2, bat RaTG13 and pangolin CoV ^2, 4, 5^. This has led to the notion that presence of an FCS in the SARS-CoV-2 ancestor was associated with successful zoonosis and pandemic transmission between humans. In support, the FCS is required for efficient proteolytic cleavage of SARS-CoV-2 spike ^2^, virus infection of human airway cells ^1, 2, 6, 7^, cell-cell fusion and syncytia formation ^2, 8, 9^, and transmission ^6, 7^.

Following the identification of the first SARS-CoV-2 strain circulating in humans (Wuhan-Hu-1), several variants of concern (VOCs) have emerged, each containing a constellation of mutations and further adaptations to host that have been associated with increased transmission. These major previous and current VOCs are designated Alpha (PANGO lineage B.1.1.7), Beta (B.1.351), Gamma (P1), Delta (B.1.617.2) and latterly Omicron (B.1.1.529). Of these, Alpha, Delta and Omicron have been the most successful globally, each rapidly replacing the previous dominating VOC over time (Omicron>Delta>Alpha). These VOCs have an increasing number of non-synonymous mutations in spike, which alter entry efficiency and kinetics ^10^ and enhance immune escape, including innate immunity ^11, 12, 13^. Much focus has been on spike mutations arising from selective pressure for antibody escape, however SARS-CoV-2 spike continuously adapts in other ways to the human host. For example, dominant VOCs (Alpha, Delta and Omicron) harbour mutations near and within the FCS which enhances spike cleavage, indicative of evolution towards an optimised FCS ^14, 15^.

Successful viral replication and transmission requires evasion or antagonism of host defensive processes, notably innate immunity, and is particularly important for zoonotic viruses that must adapt quickly or be suitably pre-adapted to the new host. Innate immune activation upregulates host cell proteins termed restriction factors that target key steps in viral replication to limit and control infection. Guanylate binding proteins (GBP) are type-1 and type-2 interferon-stimulated genes (ISGs) and a subfamily of guanosine triphosphatases (GTPases) that can act as intracellular antiviral restriction factors ^16^. GBP2 and 5 potently inhibit furin-mediated processing of viral envelope proteins, inhibiting infection of HIV-1, Influenza A Virus, Zika and measles viruses, all of which require furin cleavage for optimal infectivity ^16, 17, 18, 19^. Notably, GBPs are upregulated by type 1 and 2 interferon ^16^ and during SARS-CoV-2 infection ^20, 21^. Thus, GBPs comprise a key effector of the antiviral innate immune response that can act to limit infectious virus production during replication. Likewise, the interferon-induced transmembrane (IFITM) protein family also acts broadly to block viral entry, inhibiting viral fusion with cellular membranes, including SARS-CoV-2 ^7, 12, 22, 23, 24^.

Paradoxically, in some cases IFITM2 and/or 3 have been shown to enhance SARS-CoV-2 infection by a mechanism that remains unclear ^12, 24, 25, 26^.

Here we investigated the capacity of GBP2 and 5 to inhibit SARS-CoV-2 spike cleavage and virus infectivity, and tested whether evolution of VOCs has led to escape from GBP and IFITM restriction. We find differential sensitivity of SARS-CoV-2 spikes to both GBP and IFITMs, indicative of independent adaptation to host. Notably, while Alpha and Delta have evolved to escape restriction by GBPs, we find that Omicron is uniquely sensitivity to inhibition GBP2/5 and IFITM1, 2 and 3, consistent with Omicron displaying altered spike-mediated fusion, cell entry and tropism.

## Results

### GBP2 and 5 inhibit Wuhan-Hu-1 and Omicron, but not Alpha and Delta spike-mediated infectivity

To establish an assay for GBP restriction, we first used HIV-1, a virus whose infectivity is potently inhibited by GBP2 and 5 that block furin-mediated cleavage of the HIV-1 envelope glycoprotein ^17, 18^. As expected, exogenous expression of GBP2 and 5 in virus-producing 293T cells inhibited the infectivity of both replication-competent HIV-1, and HIV-1 Env pseudotyped virus particles (Extended data Fig.1A-C). The isoprenylation-deficient mutants of GBP2 (GBP2 C588A) and GBP5 (GBP5 C583A) which are mislocalised in the cell ^17, 18^, lost their inhibitory activity (Extended data Fig. 1B, C), as previously shown. To determine whether GBP2 and 5 have activity against SARS-CoV-2 spike, a pseudovirus (PV) assay was used in which SARS-CoV-2 spike is incorporated into lentiviral particles (herein termed PV) (Extended data Fig.1A). This allows direct comparison of how evolution of amino acid changes in spike alone (Fig. 1A) has influenced GBP sensitivity, without confounding contributions of other SARS-CoV-2 variant proteins on infectivity. 293T cells were co-transfected with plasmids encoding the spike, lentiviral genome and increasing doses of GBP-expressing plasmid. GBP expression was confirmed by flow cytometry staining for the HA-tag (Fig. 1B, Extended data Fig. 1D). Wuhan-Hu-1 PV made in the presence of GBP2 or 5 in producer cells was significantly less infectious (50%) when titrated onto Caco2 target cells, with both GBP2 and 5 inhibiting PV infectivity in a dose dependent manner (Fig. 1C). We selected naturally-permissive intestinal epithelial Caco2 cells as targets for PV infection for their endogenous expression of both ACE2 and TMPRSS2 ^27, 28^. As expected, the mutants GBP2 C588A and GBP5 C583A showed no inhibitory activity against SARS-CoV-2 infectivity (Fig. 1C). Inhibition of Wuhan-Hu-1 PV infectivity was not due to lack of spike expression on 293T cells as expressing GBPs did not alter plasma membrane levels of spike measured by flow cytometry (Extended data Fig. 2). Alpha and Delta spikes were resistant to GBP2 and 5 restriction, with no inhibition of PV infectivity (Fig. 1E, F). Strikingly, PVs containing the Omicron BA.1 (Omicron) spike were sensitive to restriction by GBP2 and 5 evidenced by a significant 50% loss of infectivity, thus behaving like Wuhan-Hu-1 (Fig.1G). Consistent with the loss of Wuhan-Hu-1 PV infectivity being mediated by GBP2/5 inhibition of furin cleavage (and not off target effects), no difference in infectivity was seen when PV were titrated onto Vero.E6 cells (Extended data Fig. 1E) that do not require furin-processing for SARS-CoV-2 infection ^2, 6^. Moreover, GBP2/5 expression significantly reduced infectivity mediated by MERS-CoV spike, which contains a furin-cleavage site ^29^, but not by SARS-CoV-1 spike that lacks a furin-cleavage site (Extended data Fig. 1F-I). Measuring GBP-restriction of SARS-CoV-2 PV infectivity on HeLa-ACE2 cells gave similar results (Extended data Fig. 1J-M), with the exception of Omicron where interestingly, PV made in the presence of GBP2/5 showed enhanced infectivity on HeLa-ACE2 cells (Extended data Fig. 1N), which we discuss later. We also noted that Omicron PV infectivity was significantly lower than other variants tested (Fig. 1H), consistent with reports that Omicron displays reduced infectivity in cell lines commonly used to study SARS-CoV-2 entry ^30, 31, 32, 33^ and in animal models ^33^. Alpha, Delta and Omicron all contain the infectivity-enhancing D614G mutation in spike (Fig. 1A) that manifested as an early host adaptation in Wuhan-like isolates and is now ubiquitous in circulating SARS-CoV-2 variants ^34, 35, 36^. Introducing the D614G mutation into Wuhan-Hu-1 spike (Wuhan-Hu-1 D614G) completely rescued PV infectivity from inhibition by GBPs (Fig. 1D). Consistent with previous reports, D614G rendered Wuhan-Hu-1 PV more infectious on a per particle basis, but remained less infectious than Alpha and Delta which also contain D614G (Fig. 1H) ^34, 35, 36^.

**Figure 1:**
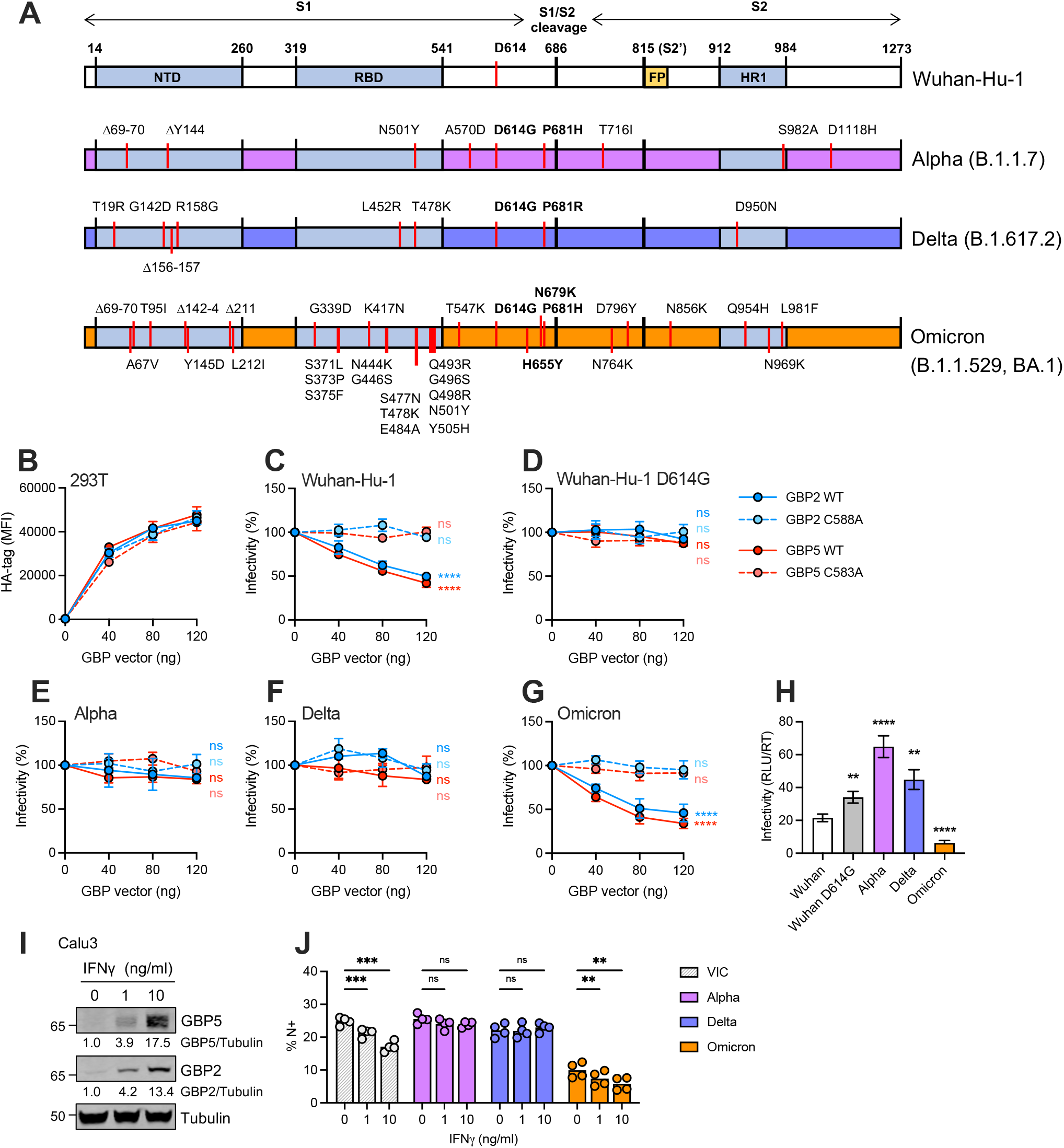
GBP2 and 5 inhibit Wuhan-Hu-1 and Omicron, but not Alpha and Delta spike-mediated infectivity. **A)** Schematic of SARS-CoV-2 spike subunit and main domain boundaries, with positions of S1/S2 and S2’ cleavage sites. Lineage defining mutations for VOCs are indicated. NTD, N-terminal domain; RBD, receptor-binding domain; FP, fusion peptide; HR1, heptad repeat 1. **B)** Expression of HA-tagged GBPs in pseudovirus PV-producing 293T cells measured by flow cytometry. **C-G)** Infectivity of PVs produced by 293T cells in the presence of increasing amounts of plasmid encoding GBP2, GBP2 C588A, GBP5 or GBP5 C583A was measured by luciferase assay (RLU) on Caco2 cells. Shown are percentage infectivity of PV made in the presence of GBPs normalised to empty vector (EV) control (no GBP, set to 100%). Percent infectivity of **C)** Wuhan-Hu-1, **D)** Wuhan-Hu-1 D614G, **E)** Alpha, **F)** Delta, and **G)** Omicron spike PV infection are shown. **H)** Comparison of particle infectivity (RLU/RT) of PVs on Caco2 cells in the absence of GBPs. Shown are the mean ±SEM from three independent experiments. **I-J)** Calu3 cells were treated with indicated doses of IFNγ for 8 h prior to infection with indicated SARS-CoV-2 variants for 36 h. **I)** Cell lysates were immunoblotted for GBP2, GBP5 and tubulin. Quantification shows relative expression of GBP2/5 over tubulin and normalised to untreated control. **J)** Equal doses (E copies/cells) of SARS-CoV-2 virus produced in cells from (I) were used to infect Caco2 cells. At 24 h, infection levels were determined by intracellular staining for nucleocapsid (N) protein. Percentage positive cells are shown (% N+ cells). Bars show the mean and individual values from two independent experiments. Two-way ANOVA (C-G, J) or repeated measured (RM) one-way ANOVA (H) with Dunnett’s post-test was used. C-G) Statistical significance for GBPs (120 ng) compared EV control is indicated. ns, not significant; *p < 0.05; **p < 0.01; ***p < 0.001; ****p < 0.0001.

Next, we sought to determine whether GBP2/5 also inhibit SARS-CoV-2 live virus infectivity. To do so, Calu3 cells were pre-treated with IFNγ, a potent inducer of GBPs ^16, 18^, to upregulate expression of GBP2/5 (Fig. 1I) and then infected with SARS-CoV-2 isolates: VIC (an early-lineage, Wuhan-like isolate) and VOCs Alpha, Delta and Omicron (Extended data Fig. 3A). Infectivity of VIC and Omicron viruses was significantly reduced by IFNγ treatment of virus producing cells (in a dose-dependent manner) evidenced by reduced infection of Caco2 target cells (Fig. 1J). By contrast, Alpha and Delta infectivity was not inhibited, consistent with data obtained using spike PV and GBP2/5 overexpression. Similar results were obtained using HeLa-ACE2 as target cells to measure infectivity (Extended data Fig. 3B).

### GBP2 and 5 inhibition of SARS-CoV-2 spike cleavage

To define mechanism, we analysed the effects of GBPs on spike S1/S2 cleavage by western blotting. Visualising uncleaved (S) and cleaved spike (S2, S2/S ratio) revealed clear differences in processing of Wuhan-Hu-1 spike on PV particles and in cell lysates in the presence of GBP2 and 5, compared to either no GBPs, or GBP2 C588A and GBP5 C583A mutants (Fig. 2A, B). Quantifying this across all spike variants showed that GBP5 significantly reduced the amount of cleaved spike (lower S2/S ratio) in Wuhan-Hu-1, Wuhan-Hu-1 D614G, Alpha, Delta and Omicron PV particles (Fig. 2C). Differential spike cleavage in the presence of GBP5 was also evident in cell lysates, showing decreased S2/S ratios (Extended data Fig. 4A), consistent with GBP perturbation of intracellular spike processing. SARS-CoV-2 viruses produced by IFNγ-treated Calu3 cells showed reduced incorporation of cleaved S2 subunit into virions (S2/N ratio) with increasing doses of IFNγ and GBP induction (Fig. 2D). Virus produced from Calu3 cells incorporated predominantly cleaved (S2) spike and low levels of uncleaved (S) spike, precluding quantification of S2/S cleavage ratios in viral lysates. While we could not detect a clear difference in S2/S ratios in virus or cell lysates (Extended data Fig. 4B) from live virus assays, the striking reduction in S2 spike incorporated into virions in the presence of GBPs is agreement with the data from PV assays.

**Figure 2:**
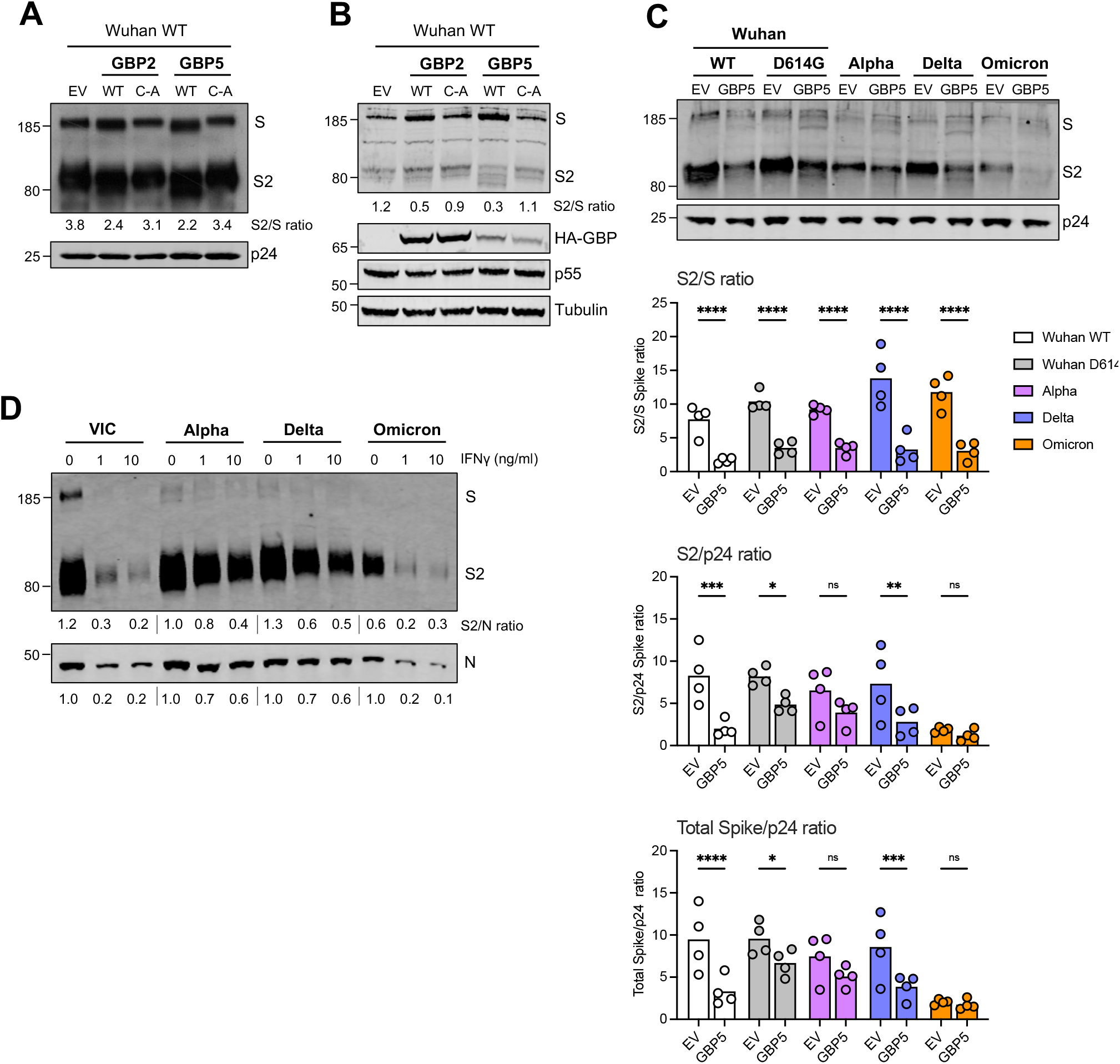
GBP2 and 5 inhibit SARS-CoV-2 spike cleavage. **A-C)** Spike PVs were produced in 293T cells in the presence of 80 ng of GBP plasmid or empty vector (EV) control. PVs and producer cell lysates were immunoblotted for spike, lentiviral Gag (p24 and p55), GBP (HA-tag) and tubulin. **A)** Immunoblot of Wuhan-Hu-1 spike from PV produced by 293T cells made in presence of GBP2 (or C588A mutant, C-A) and GBP5 (or C583A mutant, C-A) or EV control. **B)** Producer 293T cell lysate from A. **C)** Immunoblot of VOC spike PV produced in presence of GBP5 or EV. A representative immunoblot is shown. Graphs show quantification pooled from four independent experiments, measuring the proportion of cleaved spike in PV (S2/S) (top), cleaved spike incorporation (S2/p24) (middle) and total spike incorporation ((S2+S)/p24) (bottom). Mean and individual values are shown. **D)** Calu3 cells were treated with indicated doses of IFNγ for 8 h and infected with indicated SARS-CoV-2 variants for 36 h. Clarified viruses were sucrose-purified from equal volumes of culture supernatants and immunoblotted for spike (S) and nucleocapsid (N) protein. Quantification shows proportion of cleaved spike incorporation into virions (S2/N) and N intensity normalised to untreated control (0 ng/ml IFNγ). Two-way ANOVA with Dunnett’s post-test was used. ns, not significant; *p < 0.05; **p < 0.01; ***p < 0.001; ****p < 0.0001.

Next, we tested whether decreasing spike expression can sensitise Wuhan-Hu-1 D614G, Alpha and Delta PV to GBP2/5 restriction. Indeed, lowering the expression of spike by transfecting decreasing amounts of spike plasmid during PV production restored the ability of GBP2/5 to restrict Wuhan-Hu-1 D614G, and to a lesser extent Alpha and Delta infectivity to levels seen for Wuhan-Hu-1 (Extended data Fig. 5A-D), whereas Omicron spike was even further inhibited by GBP2/5 (Extended data Fig. 5E). Similar results were obtained when measuring PV infectivity on HeLa-ACE2 cells, although Omicron again behaved as an outlier (Extended data Fig. 5F-J). To determine whether increasing spike incorporation into PV could render Wuhan-Hu-1 resistant to GBP5 restriction, we took an advantage of the fact that SARS-CoV-2 spike contains an endoplasmic reticulum retention signal in the cytoplasmic tail (CT) and that truncation of the last 19 residues of the spike cytoplasmic tail (ΔCT) removes this motif, boosting spike incorporation and particle infectivity during pseudotyping ^37, 38^. Concordantly, truncation of the Wuhan-Hu-1 spike CT (Wuhan ΔCT) enhanced PV infectivity 10-fold compared to PV containing the full-length spike (Extended data Fig. 6A). However, both ΔCT and full-length spike remained sensitive to GBP5 restriction (Extended data Fig. 6B), although ΔCT spike was less sensitive compared to its full-length counterpart (30% and 50% inhibition, respectively). Immunoblotting confirmed both the increase in incorporation of ΔCT spike into PV, and GBP5 inhibition of ΔCT and full-length spike cleavage (Extended data Fig. 6C). Taken together these data identify GBP2 and 5 as restriction factors that target SARS-CoV-2 by interfering with spike processing at the S1/S2 cleavage site. Consequently, the infectivity of early-lineage variants (Wuhan-Hu-1/VIC) is inhibited, while VOCs Alpha and Delta have evolved to escape this inhibition, whereas Omicron has not.

### Effects of GBP 2 and 5 on spike-mediated cell entry routes

In order to test whether GBP inhibition of spike cleavage influences the entry route into target cells, we first confirmed the entry phenotype on Caco2 cells using inhibitors Camostat (a serine protease inhibitor that blocks TMPRSS2) and E64d (an inhibitor of endolysosomal cathepsins) (Fig. 3). Early-lineage SARS-CoV-2, and VOCs Alpha and Delta are inhibited by Camostat, but not E64d in cells expressing TMPRSS2, a finding we confirm here using Caco2 cells (Fig. 3A-D) ^22, 28, 39^. By contrast, Omicron was potently inhibited by E64d, and was less sensitive than other SARS-CoV-2 variants to Camostat in both PV and live virus infection assays (Fig. 3A-D), suggesting Omicron more efficiently uses TMPRSS2-independent endosomal entry pathway, concordant with recent studies ^30, 31, 32, 33, 40^. Furin-cleavage at spike S1/S2 can influence viral entry route. Therefore, we directly compared Omicron spike cleavage to other VOCs, and early-lineage isolates VIC and IC19, by immunoblotting viral particles purified from infected Caco2 culture supernatants. Evolution of VOCs for an optimised FCS correlated with cleavage efficiency, with Delta>Alpha>IC19/VIC (Fig. 3E). Notably, Omicron virus incorporated predominantly cleaved spike, with little or no uncleaved spike detected in virions (Fig. 3E), a result confirmed using two independent Omicron isolates. The total amount of cleaved spike in virions (S2/N) was, however, lower for Omicron compared to other VOCs (Fig. 3E). Efficient S1/S2 cleavage of Omicron spike is consistent with this VOC containing the FCS optimising mutations P681H, N679K and H655Y (Fig. 1A) that can enhance S1/S2 cleavage ^14, 15^. These data suggest that Omicron’s shift in cellular tropism is unlikely to be explained by inefficient spike cleavage in virions.

**Figure 3:**
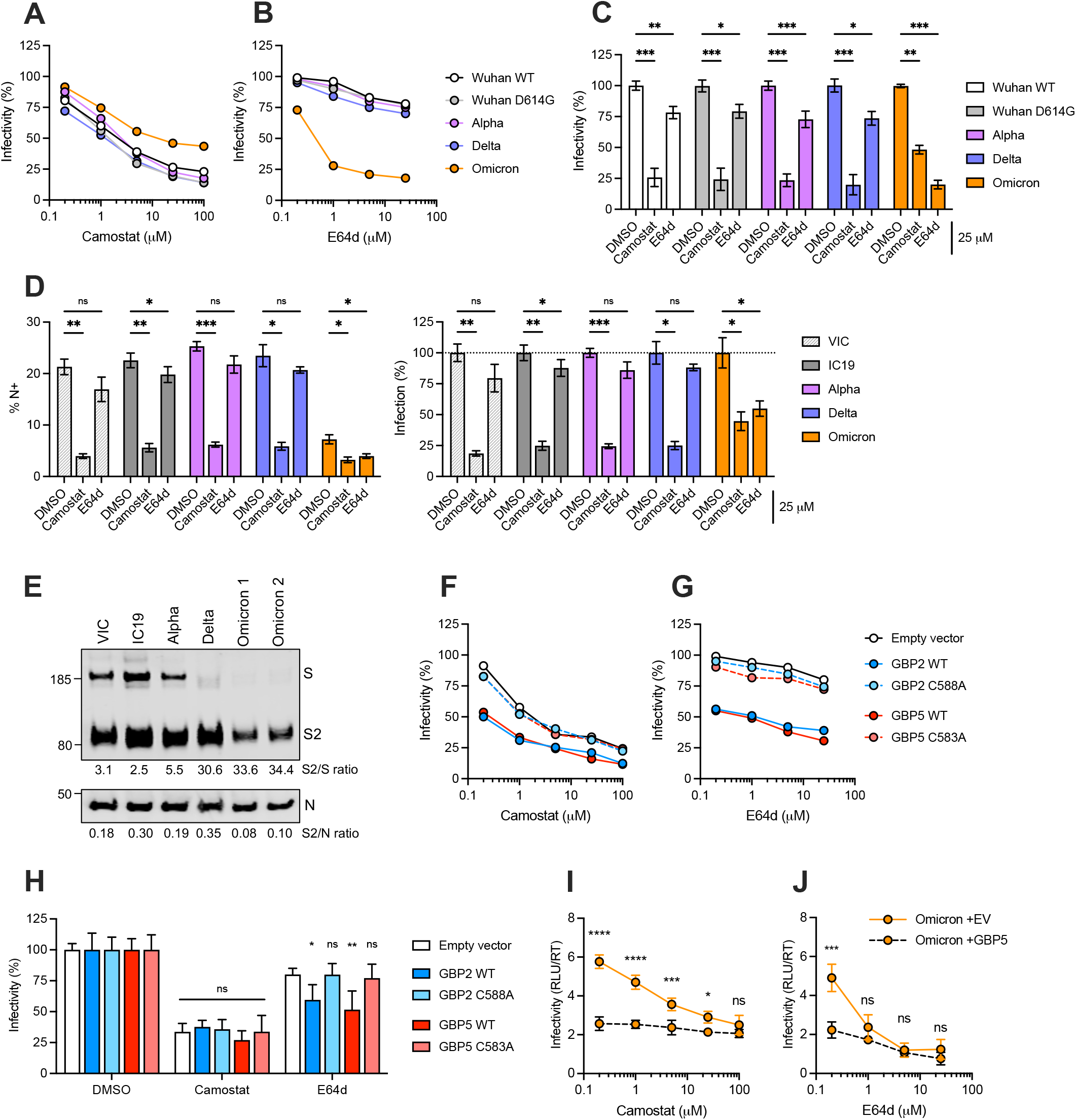
Effects of GBPs on spike-mediated entry pathways in Caco2 cells. Inhibition of spike-mediated PV infection of Caco2 cells by **A)** Camostat and **B)** E64d inhibitors. Shown is a representative titration. **C)** Inhibition of PV infection of Caco2 cells in the presence of Camostat or E64d normalised to DMSO control. Infection was measured by luciferase assay. **D)** Inhibition of SARS-CoV-2 live virus infection of Caco2 cells by Camostat or E64d. Infection was measured by at 24 h by intracellular flow cytometry staining for nucleocapsid (N) protein. Percentage N+ cells (% N+) (left) and percentage infection for each virus normalised to the corresponding DMSO control (right) are shown. **E)** Immunoblot analysis of spike cleavage in SARS-CoV-2 virions. Quantification shows relative proportion of cleaved spike (S2/S) and cleaved spike incorporation into virions (S2/N). Two independent Omicron isolates are shown. **F-H)** Effect of GBPs on SARS-CoV-2 spike-mediated entry pathways. Wuhan-Hu-1 spike PV made in presence of 80 ng of indicated GBP plasmid or EV control was used to infect Caco2 cells in the presence of increasing concentrations of **F)** Camostat or **G)** E64d. Percentage infection compared to DMSO EV control are shown. **H)** Inhibition of Wuhan-Hu-1 PV +/- GBPs in the presence of 25 μM Camostat or E64d. Data are normalised to DMSO control separately for each GBP treatment and grouped by inhibitor treatment. **I-J)** Omicron spike PV was made in presence of GBP5 or EV control and infection of Caco2 cells in the presence of increasing doses of **I)** Camostat or **J)** E64d. Mean ±SEM from three independent experiments are shown. Two-way ANOVA with Dunnett’s post-test was used. ns, not

Measuring the sensitivity of PV made in the presence of GBPs to entry inhibitors showed that GBP2 and 5 partially, but significantly, sensitised Wuhan-Hu-1 PV to inhibition by E64d, but not Camostat, on Caco2 cells (Fig. 3F-H). For Omicron, addition of E64d, but not Camostat, further reduced infection of PV made in the presence of GBP5 (Fig. 3I, J). Thus although GBP-mediated inhibition of spike cleavage leads in somewhat increased endosomal entry, it does not make PV completely TMPRSS2-independent. Of note, HeLa-ACE2 cells support more endocytic entry of SARS-CoV-2, evidenced by E64d, but not Camostat, inhibiting infection of IC19, VIC, Alpha and Delta viruses in these cells (Fig. S7).

This likely explains the observation that Omicron PV made in the presence of GBP5 shows enhanced infectivity of HeLa-ACE2 cells, and is not less infectious on 293T-ACE2 cells (Extended data Fig. 1 and 8), reflecting Omicron’s preference for endosomal entry.

### GBPs do not sensitise to IFITM restriction

Having shown that GBPs perturb furin cleavage of spike, partially redirecting PV towards endosomal entry, we tested whether GBPs sensitise PV to inhibition by endosomal IFITM2 and 3. IFITMs can target SARS-CoV-2 spike fusion with host cell membranes to inhibit infection ^7, 12, 22, 23, 24, 25^ and are differentially localised in cells, with IFITM1 being found mostly at the plasma membrane and IFITM2/3 predominantly endosomal. To explore the relationship between these two innate immune restriction factors, we used Caco2 cells stably over-expressing either IFITM1, 2 or 3 (Fig. 4A) and confirmed the expected IFITM localisation by immunofluorescence microscopy (Fig. 4B). We first established the IFITM restriction phenotype in these cells and consistent with TMPRSS2-dependent plasma membrane entry, IFITM1 was found to potently and significantly inhibit infection of early-lineage viruses (VIC, IC19) and VOCs Alpha and Delta (Fig. 4C-F). By contrast, IFITM2 and IFITM3 did not inhibit, but rather enhanced infection (Fig. 4C-F), in agreement with previous studies ^12, 22, 24, 25, 26^. Similar results were obtained using PV infection assays (Fig. 4H-K).

**Figure 4:**
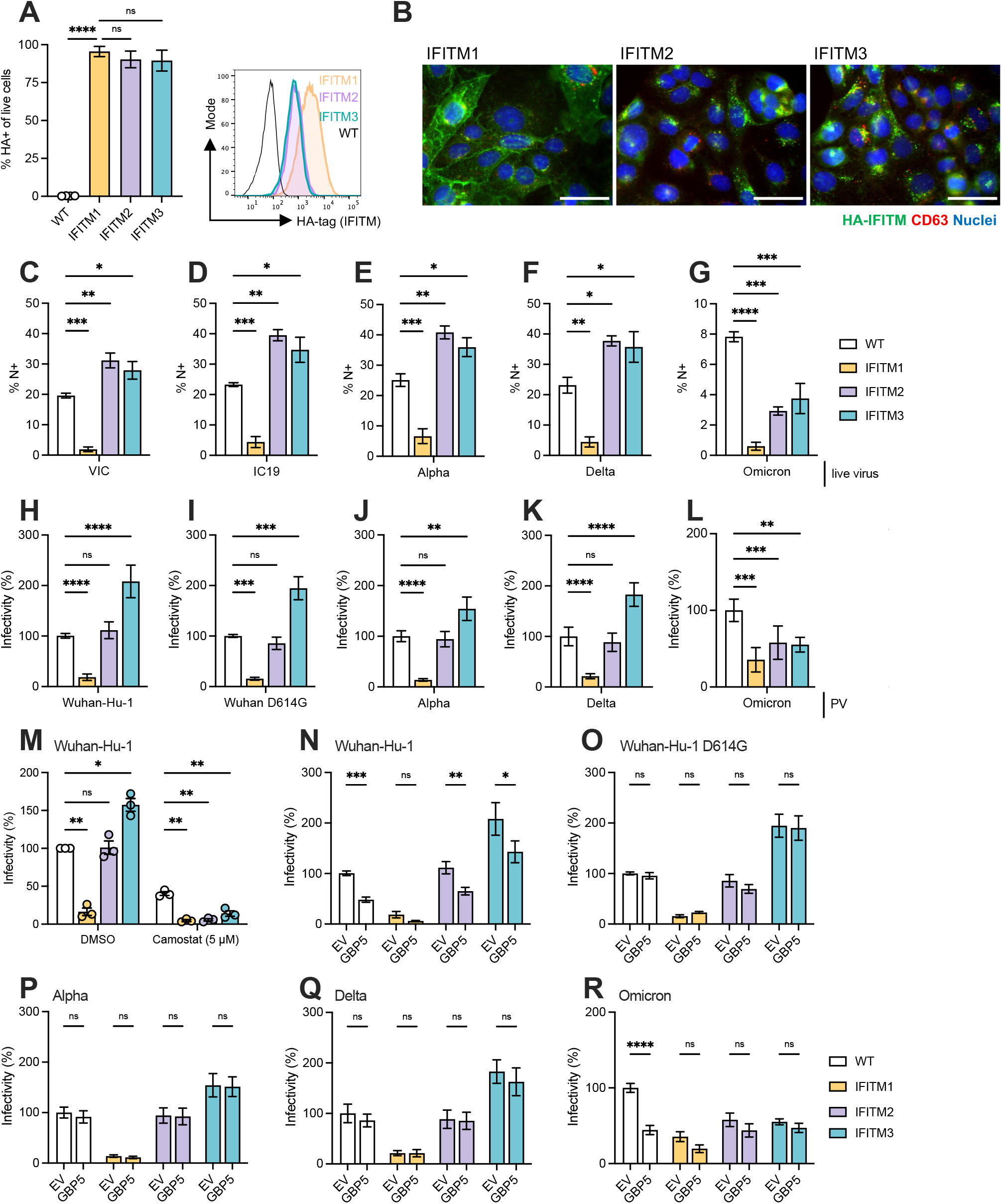
IFITM restriction of SARS-CoV-2 and the interplay with GBPs. A)Stable expression of HA-tagged IFITMs 1-3 in Caco2 cells measured by flow cytometry. Representative histogram is shown (right). **B)** Immunofluorescence images of IFITM1-3 expression in cells from A). HA-tag IFITM (green), CD63 (red), and nuclei shown in blue. Scale bar is 50 μm. **C-G)** IFITM-expressing Caco2 cells were infected with live SARS-CoV-2. Shown is percent N+ cells at 24 hpi of **C)** VIC, **D)** IC19, **E)** Alpha, **F)** Delta, and **G)** Omicron isolates. **H-L)** PV infection of IFITM-expressing Caco2 cells with **H)** Wuhan-Hu-1, **I)** Wuhan-Hu-1 D614G, **J)** Alpha, **K)** Delta, and **L)** Omicron spike PV. Percent infectivity normalised to WT Caco2 cells (no IFITM over-expression) are shown. **M)** IFITM-expressing Caco2 cells were infected with Wuhan-Hu-1 PV in the presence of 5 μM Camostat or DMSO control. Percent infectivity normalised to DMSO-treated WT Caco2 cells is shown. **N-R)** IFITM-expressing Caco2 cells were infected with spike PVs made in presence of GBP5 or EV control. Infectivity normalised to EV control on WT cells for PV made using **N)** Wuhan-Hu-1, **O)** Wuhan-Hu-1 D614G, **P)** Alpha, **Q)** Delta, and **R)** Omicron spikes is shown. Mean ±SEM from three independent experiments are shown. One-way ANOVA (A-L) or two-way ANOVA (M-R) with Dunnett’s post-test were used. ns, not

Blocking TMPRSS2-mediated entry with Camostat sensitised Wuhan-Hu-1 PV to restriction by endosomal IFITM2 and 3, leading to an almost complete inhibition of infection (Fig. 4M). Testing the effects of GBP5 expressed in PV producer cells, on the sensitivity of PV to IFITM restriction in target cells, we found no significant difference in IFITM1 inhibition of Wuhan-Hu-1 PV made in the presence or absence of GBP5 (Fig. 4N). For IFITM2/3, although PV made in the presence of GBP5 appeared to show enhanced sensitivity to endosomal IFITM2/3, this can be accounted for by the inhibitory effect of GBP5, evidenced by comparing infectivity of GBP5 plus IFITM2 or 3, to GBP5 alone (Fig. 4N). Moreover, Wuhan-Hu-1 D614G, Alpha and Delta PV sensitivity to IFITMs was similarly unaffected by GBPs (Fig. 4O-Q). Given that GBPs did not completely prevent S1/S2 spike cleavage in our pseudovirus assay, this result is not unexpected. Similar results were obtained using live viruses (VIC, Alpha and Delta) produced in presence of IFNγ, where inducing expression of GBP2/5 did not sensitise virus to enhanced IFITM restriction (Fig. S9). Collectively these data show that while combining GBPs and IFITMs can lead to an overall potent reduction of infection, inhibiting furin cleavage by GBPs does not make a virus that is resistant to IFITMs become sensitive to restriction.

### Omicron virus is restricted by IFITM 1, 2 and 3

The recently emerged Omicron variant has acquired over 30 spike mutations compared to Wuhan-Hu-1 (Fig. 1A). Having shown that Omicron was sensitive to GBP inhibition, we tested Omicron’s sensitivity to IFITMs. Strikingly, we found Omicron was unique amongst SARS-CoV-2 viruses we tested, being sensitive to inhibition of infection by IFITM1, 2 and 3 in Caco2 cells (Fig. 4G). Similar restriction of Omicron by IFITM1, 2 and 3 was seen in PV assays (Fig. 4L). Next, we investigated the effects of combining GBP and IFITMs on Omicron infection. Fig. 4R shows that Omicron PV made in the presence of GBP5 was not significantly more sensitive to IFITM restriction. Similarly, IFNγ treatment in producer cells did not further sensitise Omicron to IFITM inhibition (Extended data Fig. 9D) To test whether increased spike incorporation changes Omicron sensitivity to GBP or IFITM restriction, we used full-length and ΔCT spike mutants (described in Extended data Fig. 6). As expected, increasing spike incorporation into Omicron PV by truncation of the spike CT (Omicron ΔCT) enhanced PV infectivity 70-fold compared to PV containing the full-length Omicron spike. (Fig. 5A). However, there was no difference in GBP5 restriction of Omicron PV containing ΔCT spike compared to full-length spike (Fig. 5B), and no rescue from restriction by IFITMs (Fig. 5D-F). Enhanced spike incorporation of the ΔCT mutant and the GBP5 induced spike cleavage defect were confirmed by immunoblotting (Fig. 5C). Omicron BA.1 and BA.2 isolates have similar PV particle infectivity and were similarly restricted by GBP5 (50% and 42% inhibition, respectively) (Extended data Fig. 10). Moreover, both BA.1 and BA.2 PV are more sensitive to inhibition by E64d compared to Camostat and show similar restriction by IFITM1-3 (Extended data Fig. 10B-D). Taken together, these data reveal striking differences between Omicron and preceding SARS-CoV-2 isolates in sensitivity IFITM restriction, indicative of an altered Omicron entry route.

**Figure 5:**
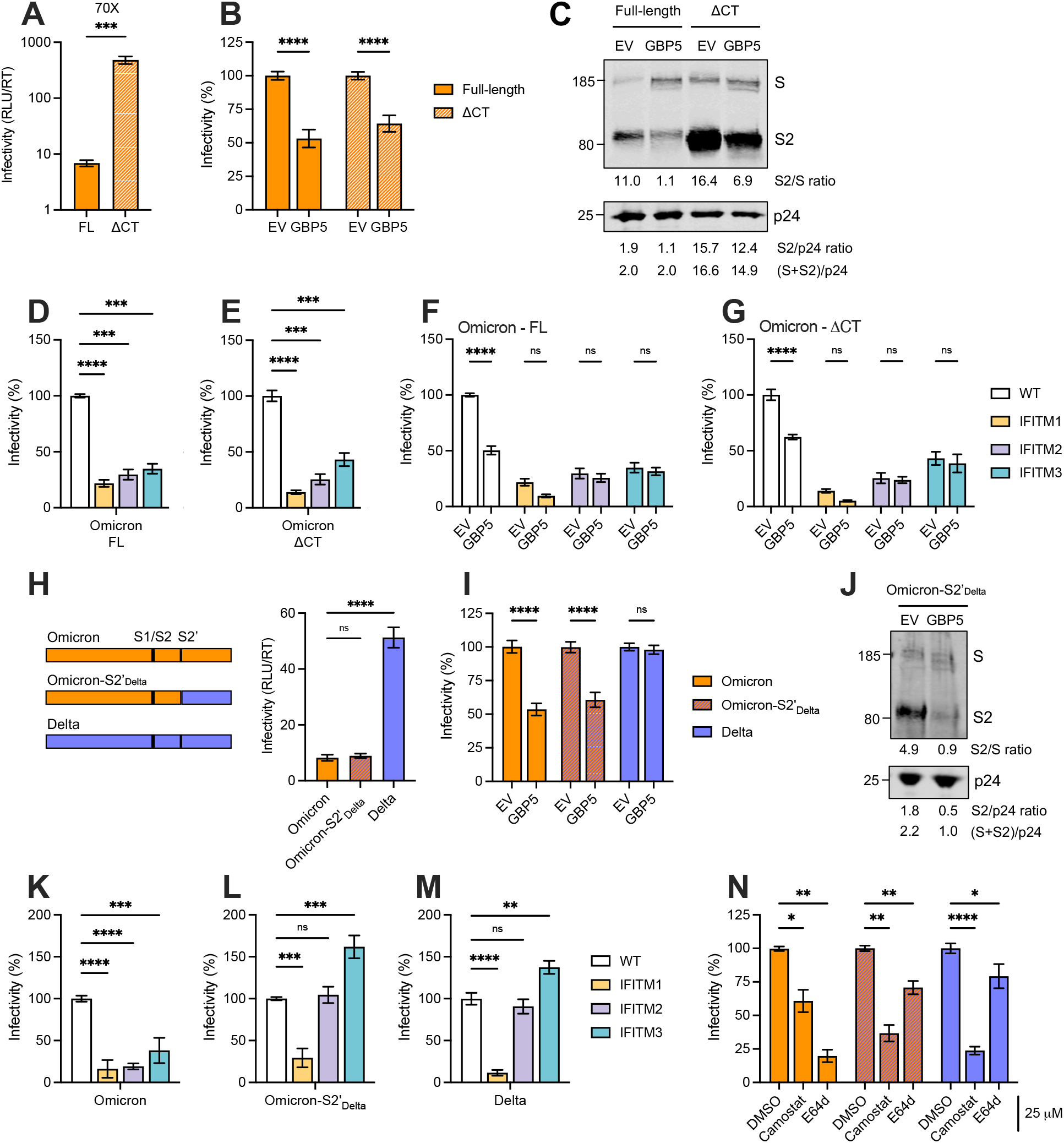
Effects of spike C-terminal truncation and the S2 domain on Omicron sensitivity to GBP and IFITMs and entry route. **A-C)** Omicron spike PV were produced in presence of GBP5 or EV control using plasmids encoding either full-length (FL) or a 19 residue C-terminal truncation (ΔCT) Omicron spike. Spike PV were titrated on WT Caco2 cells, showing **A)** raw infectivity (RLU/RT) values in absence of GBP5 expression (EV control) and **B)** percent infectivity normalised to EV control for each Omicron spike (FL and ΔCT). **C)** Omicron FL and ΔCT spike PVs were produced in the presence of GBP5 or EV control and immunoblotted for spike and lentiviral Gag (p24). Quantification shows relative proportion of cleaved spike in PV (S2/S), cleaved spike incorporation into PV (S2/p24) and total spike incorporation ((S2+S)/p24). **D-G)** Indicated Omicron PV were titrated on Caco2 WT and IFITM expressing cells. Shown is infectivity normalised to EV control for **D)** FL Omicron PV, **E)** ΔCT Omicron PV, **F)** FL Omicron PV made in presence of GBP5 or EV control and **G)** ΔCT Omicron PV made in presence of GBP5 or EV control. **H-N)** Omicron spike S2’ domain was swapped with Delta S2’ domain to produce Omicron-S2’_Delta_ spike (schematic). **H-I)** Omicron, Omicron-S2’_Delta_, and Delta spike PV were produced in presence of GBP5 or EV control and titrated on WT Caco2 cells, showing **H)** raw infectivity (RLU/RT) values in absence of GBP5 expression (EV control) and **I)** percent infectivity normalised to EV control for each spike PV. **J)** Omicron-S2’_Delta_ spike PV made in presence of GBP5 or EV control were immunoblotted for spike and p24. **K-M)** PV infection of IFITM transduced Caco2 cells with **K)** Omicron, **L)** Omicron-S2’_Delta_, and **M)** Delta spike PV. Percent infectivity normalised to WT Caco2 cells (no IFITM over-expression) are shown. **N)** Inhibition of spike PV infection of Caco2 cells in the presence of 25 μM Camostat or E64d normalised to DMSO control. Mean ±SEM from three independent experiments are shown. Two-tailed paired t test (A, B, I); One-way ANOVA (D, E, H, L, N) or two-way ANOVA (F, G, K) with Dunnett’s post-

### Omicron S2’ domain is the determinant for altered entry route and IFITM sensitivity

Omicron (BA.1) contains 3 unique mutations (Q954H, N969K and L981F) within the heptad repeat domain 1 (HR1) of spike that mediates viral fusion, as well as N856K adjacent to the fusion peptide (Fig. 1A), which we hypothesised may play a role in Omicron’s altered entry and sensitivity to spike targeting restriction factors. Since Delta spike has the highest fusogenicity ^41^ we swapped S2’ domain of Omicron spike with that of Delta to produce pseudoparticles with Omicron-S2’_Delta_ spike chimera (Fig. 5H). PV containing the chimeric Omicron-S2’_Delta_ spike PV retained low particle infectivity (Fig. 5H), and remained sensitive to GBP5 inhibition of spike cleavage, behaving like WT Omicron and not like Delta (Fig. 5I-J).

However, replacing the S2’ domain of Omicron with that of Delta, rendered Omicron-S2’_Delta_ spike PV resistant to IFITM2 and 3 inhibition, and sensitive to IFITM1, thus behaving like Delta (Fig. 5K-M). To test whether Omicron-S2’_Delta_ spike’s resistance to endosomal IFITM2/3 was due to altered cell entry we compared sensitivity to inhibitors and found that Omicron-S2’_Delta_ PV was sensitive to the TMPRSS2 inhibitor Camostat and resistant to the endosomal E64d inhibitor, thus behaving like Delta (Fig. 5N). Taken together, these data show that residues within the S2’ domain of spike are the determinant for IFITM sensitivity by modulating entry route, notably regulating Omicron’s use of TMPRSS2-independent, endosomal pathways.

## Discussion

Innate immunity is a potent first-line host cell defense against viruses, upregulating a group of interferon-stimulated genes (ISGs) which can act directly as restriction factors, targeting key steps in viral replication and collectively inducing an antiviral state. SARS-CoV-2 triggers innate immune sensing and induces an interferon-response ^11, 27, 42^, upregulating canonical ISGs including GBP2 and 5 in primary human airway epithelial cells ^21^. Evolution of mutations outside of spike allow for SARS-CoV-2 evasion/antagonism of innate immune sensing ^11^; however, spike itself can be targeted by innate immune responses. Here we discover that the interferon-inducible restriction factors GBP2 and 5 interfere with furin-mediated cleavage of SARS-CoV-2 spike and potently inhibit infection by the early-lineage SARS-CoV-2 strain Wuhan-Hu-1 but that dominant VOCs have evolved differential sensitivity to restriction. Using spikes from other coronaviruses with pandemic potential, MERS-CoV and SARS-CoV-1, we confirm that restriction by GBP2/5 correlates with the requirement for furin-mediated spike processing, such that MERS-CoV spike is sensitive, but SARS-CoV-1 resistant, to inhibition by GBPs.

Alpha and Delta have both evolved an optimised FCS by acquiring P681H and P681R respectively, leading to enhanced S1/S2 spike cleavage by furin ^12, 14, 15^, with other mutations outside the FCS (i.e. H69/V70 deletion and D614G) also influencing spike cleavage ^15, 43, 44^. Therefore, we hypothesised that these VOCs may have evolved to evade GBP restriction.

Indeed, we found that both Alpha and Delta were resistant to GBP-mediated inhibition of PV and live virus infectivity. However, escape from restriction by these VOCs cannot be simply attributed to an optimised FCS overcoming GBP inhibition of spike cleavage. This is because both Alpha and Delta displayed the same GBP-mediated spike cleavage defect as Wuhan-Hu-1. Moreover, the amount of cleaved spike incorporated into PV was comparable between Wuhan-Hu-1 (restricted) and Alpha and Delta (not-restricted). In addition, Omicron PV and virus infectivity was potently inhibited by GBP2/5, despite containing similar FCS-optimising mutations as Alpha/Delta. Instead, we argue that evolution of Alpha and Delta for enhanced activity of spike, including better fusogenicity and infectivity ^6, 7, 9, 41, 45^, are likely key determinants mediating their escape from restriction. In support, we find a correlation between inherent infectivity and the GBP restriction phenotype, such that Alpha and Delta which are more infectious on Caco2 cells are not restricted by GBPs, whereas less infectious Wuhan-Hu-1 and Omicron are restricted ^9, 30, 31, 45^. Consistent with this notion, Wuhan-Hu-1 D614G mutant spike that has enhanced infectivity was rescued from GBP inhibition. Therefore, we propose that it is the combined effects of mutations that have accumulated in Alpha and Delta spikes, changing spike activity and infectivity, that have led to evolution of these VOCs to escape inhibition by GBPs.

IFITMs are another family of ISGs that can target SARS-CoV-2 spike and restrict infection, in this case by inhibiting fusion of viral and cellular membranes in target cells, therefore we sought to investigate the relationship between GBP and IFITMs. Consistent with TMPRSS2-dependent plasma membrane entry of Wuhan-Hu-1/VIC/IC19, Alpha and Delta viruses, IFITM1, but not IFITM2 and 3, inhibited infection of natural-permissive Caco2 cells. As expected, inhibiting plasma membrane entry with Camostat and redirecting entry towards a more endosomal pathway led to restriction of Wuhan-Hu-1 by IFITM2 and 3. However, although perturbing furin cleavage with GBPs increased sensitivity to E64d (indicative of increased endosomal entry), GBPs did not further sensitise Wuhan-Hu-1 PV to inhibition by endosomal IFITM2 or 3. This is in contrast to studies showing that interfering with spike cleavage by mutating the FCS in Alpha sensitises virus to inhibition by endosomal IFITM2 ^12, 22^. We explain this by GBPs reducing, but not completely preventing, spike S1/S2 cleavage in our PV assay, allowing for a significant proportion of PV particles to use TMPRSS2-dependent plasma membrane fusion and avoid endosomal IFITMs. That GBPs did not completely redirect SARS-CoV-2 towards endosomal entry routes is consistent with other cellular proteases mediating spike processing at the polybasic cleavage site in the absence of furin ^8^. It is notable however, that although GBPs did not sensitise to IFITMs, combining GBP5 in producer cells with IFITM1 in target cells led to an almost complete inhibition of Wuhan-Hu-1 infection, when compared to infection in the absence of these restriction factors. Thus, during an innate immune response to viral infection where multiple ISGs including GBP2/5 and IFITM1/2/3 are induced, we might expect to see stronger inhibitory effects.

Our most striking observation is that Omicron is unique amongst the VOCs we tested in being sensitive to inhibition by GBP2/5, and also IFITM1, 2 and 3. Omicron is also unique amongst SARS-CoV-2 in its entry pathway, having evolving towards TMPRSS2-independence and endosomal-dependent fusion ^30, 31, 32^. In agreement with these recent reports, we show Omicron is significantly less infectious than other SARS-CoV-2 isolates on TMPRSS2 expressing Caco2 cells and less sensitive to the TMPRSS2 inhibitor Camostat, while being more sensitive to the cathepsin inhibitor E64d, and now map this entry phenotype to the S2’ domain of spike. Omicron’s use of an endosomal entry pathway is consistent with our observation that, unlike other VOCs, Omicron is sensitive to inhibition by IFITM2 and 3, which are predominantly located within endosomes. Omicron contains the same P681H FCS-optimising mutation as Alpha, as well as addition H655Y and N679K that may further optimise the FCS ^14, 15^. Indeed, we show that Omicron S1/S2 cleavage is notably more efficient that either Alpha or Delta, with Omicron virus incorporating very little, if any, uncleaved spike into virions. Therefore, we expected that Omicron would also be resistant to GBPs, like other VOCs; however, Omicron was strikingly sensitive GBP2/5 with infectivity inhibited to levels seen for Wuhan-Hu-1. Moreover, we were unable to rescue Omicron from GBP or IFITM restriction by increasing the amount of incorporated spike into PVs, to a level that increased the baseline infectivity by 70-fold. We therefore conclude that it is the inherent differences Omicron’s spike activity, rather than differences in S1/S2 cleavage efficiency or spike incorporation, that make this VOC sensitive to GBP-mediated restriction of infection.

Omicron spike is reportedly less fusogenic ^30, 31, 32^ and compared to Wuhan-Hu-1, Omicron contains 4 unique mutations in the S2’ domain that mediates viral fusion (Fig. 1A) ^30, 46^.

These substitutions, not seen Alpha and Delta, may be factors contributing to the differences seen between Omicron and other VOCs in both restriction phenotypes and tropism. Indeed, replacing the S2’ domain in Omicron spike with that of Delta (Omicron-S2’_Delta_), converted the phenotype and rendered PV “Delta-like” and resistant to IFITM2/3 by switching Omicron PV entry back to TMPRSS2-dependence and away from endosomal entry. Interestingly, this PV retained low infectivity and sensitivity to GBP restriction, suggesting that other differences in the Omicron spike influence GBP sensitivity. Given the significant number of substitutions in Omicron, it is unsurprising the situation is complex and likely context-dependent with other changes outside of S2, perhaps in the RBD or NTD for example, potentially contributing to the Omicron phenotype. The *in vivo* interplay between Omicron and innate immunity, restriction factors and tropism remain unknown. It is possible that the expression of different innate immune restriction factors varies across cells/tissues, such that Omicron has evolved into a niche that allows it to avoid or tolerate GBPs and IFITMs.

Moreover, IFITM2 and 3 are suggested to act as co-factors for SARS-CoV-2 in some cases ^26^, and we and others also see enhancement of early-lineage isolates and Alpha and Delta infection by IFITM2/3 ^12, 22, 24, 25, 26^, so it cannot be excluded that Omicron may similarly exploit IFITMs in some settings. It is clear that SARS-CoV-2 needs to balance efficient cell entry with evasion of compartmentalised restriction factors, and it will be intriguing to see how Omicron does this to successfully infect target cells *in vivo*. Omicron has evidently evolved to do things differently, but effectively, and exploit a different cellular niche, one to which it is clearly well-adapted.

SARS-CoV-2 VOCs have evolved separately from early lineage strains and not from each other. It is therefore not surprising that these VOCs have explored different evolutionary solutions to the problems they faced, and that the selective pressures encountered by each VOC are not identical. For example, Alpha and Delta evolved prior to significant levels of adaptive immunity in the population, and so these VOCs may have faced stronger pressure to evolve spikes that evade innate immunity, including restriction factors like GBPs and IFITMs, while being less influenced by the need for humoral immune escape. By contrast Omicron, the first real antibody escape variant, evolved at a time of much greater population level humoral immunity, therefore Omicron has been exposed to different selective pressures, requiring different evolutionary solutions. Our data showing differential sensitivity of Alpha/Delta vs Omicron to GBP and IFITM restriction, as well as others showing significant Omicron antibody escape ^30, 32^, are consistent with this, reinforcing the notion that Omicron has taken a different evolutionary path to preceding VOCs. We propose a scenario in which evolution of Omicron spike for neutralising antibody escape has influenced the ability to evade innate immunity. The critical balance between viral evasion of innate and adaptive immunity has precedent. This is borne out of studies of HIV-1 evolution in a host, where HIV-1 isolates from early in infection (so called transmitter/founder viruses) are completely resistant to IFITM restriction, but overtime, the selective pressure from adaptive immunity, and the resulting neutralising antibody escape mutations in HIV-1 Env, leads to viral isolates having increased sensitivity to IFITMs and interferons ^47^. We propose that similar processes have occurred during SARS-CoV-2 evolution to host, in which the need to escape from neutralising antibody became the dominant selective pressure on Omicron, resulting in a compensatory, but tolerable, increase in sensitivity to innate immunity, while also impacting on spike activity and cell tropism. We predict that this interplay between evasion of innate and adaptive immunity, and the consequences for transmission and tropism, will be features of future SARS-CoV-2 evolution, and emergence of new VOCs, and that linking this evolution to phenotype will become important aspects for understanding and predicting SARS-CoV-2 biology, and ultimately pathogenesis.

## Materials and methods Cells

HEK293T/17 cells (abbreviated herein as 293T cells) were obtained from American Type Culture Collection (ATCC, CRL-11268). Caco2 cells were a gift from Dalan Bailey (Pirbright Institute) and originally obtained from ATCC. Calu3 cells were purchased from AddexBio (C0016001). HeLa-ACE2 (stable cell line expressing ACE2) were provided by J.E. Voss and D. Huang (Scripps Research Institute) ^48^. Vero.E6 cells were obtained from the National Institute for Biological Standards and Control (NIBSC). Vero.E6-TMPRSS2 cells (stable cell line expressing TMPRSS2 under G418 selection) were a gift from Mala Maini (UCL). 293T-ACE2 (stable cell line expressing ACE2 under puromycin selection) were a gift from Wendy Barclay (Imperial College London). HeLa-TZM-bl cells (expressing luciferase and beta-galactosidase under the control of HIV-1 LTR) were obtained from the Centre for AIDS Reagents (CFAR). All cell lines were grown in Dulbecco’s modified Eagle’s medium (DMEM, Thermo Fisher Scientific) supplemented with 10% fetal bovine serum (Labtech) and 1% Pen Strep (penicillin-streptomycin, Thermo Fisher Scientific), and maintained in humidified 5% CO2 incubators at 37°C. Cells were passaged every 2-4 days when they reached 80-90% confluency. Caco2 cells were transduced with IFITM1/2/3 lentivectors as described previously ^49^ and selected with 10 μg/ml puromycin (Merck) to produce stable cell lines expressing individual HA-tagged IFITM proteins. IFITM expression was confirmed by flow cytometry.

### Plasmids

SARS-CoV-2 Spike expression vectors were originally synthesised by Genewiz and subcloned into pcDNA3.1+ vector. All spike sequences are full-length, unless otherwise stated. Wuhan-Hu-1 WT ^50^, Wuhan-Hu-1 D614G and Alpha ^51^ spike expression vectors were gifts from Laura McCoy (UCL). Delta and Omicron BA.1 spike expression vectors were a gift from Katie Doores (King’s College London). Wuhan-Hu-1 WT, Omicron BA.1 and BA.2 spike ΔCT expression vectors were a gift from Wendy Barclay (Imperial College London). SARS-CoV-1 Spike and MERS-CoV Spike expression vectors were a gift from Joe Grove (Centre for Virus Research, Glasgow). Plasmid encoding HIV-1 Env pSVIII_JRFL was a gift from Laura McCoy (UCL). Plasmid encoding full length HIV-1 pNL4.3 was donated by Dr M Martin and obtained from CFAR. Lentiviral backbone packaging plasmid (expressing HIV Gag, Pol, Tat and Rev) p8.91 and the reporter plasmid encoding luciferase gene pCSLW were a gift from Greg Towers (UCL). GBP expression vectors encoding HA-tagged GBPs (GBP2 WT, GBP2 C588A, GBP5 WT, GBP5 C583A) and BFP reporter expressed from an IRES ^17, 18^ were a gift from Daniel Sauter (Ulm University Medical Center, Germany).

### Mutagenesis

Omicron BA.2 spike ΔCT vector was converted to full-length by mutating stop codon at position 1255 into lysine residue (*1255K) as found in WT sequence. Mutagenesis was performed using QuikChange Lightning site-directed mutagenesis kit (Agilent) and following primers: 5’-GTCCTCGTCGAACTTGCAGCAGCTGCCAC-3’ and 5’-GTGGCAGCTGCTGCAAGTTCGACGAGGAC-3’.

Omicron chimera was constructed by Gibson assembly cloning method. Delta S2’ domain sequence was PCR amplified using following primers:

5’-GCAGTATGGCGATTGTCTGGGCGAC-3’

5’-CGTACACGGTATTGTTCACAATGCCGATCAC-3’.

Omicron BA.1 vector sequence excluding S2’ domain was amplified using following primers: 5’-GTGATCGGCATTGTGAACAATACCGTGTACG-3’

5’-GTCGCCCAGACAATCGCCATACTGC-3’

The fragments were assembled using NEBuilder HiFi DNA Assembly Cloning Kit (New England Biolabs) according to manufacturer’s instructions. All mutagenesis and cloning was confirmed by sequencing.

### SARS-CoV-2 viruses

SARS-CoV-2 isolates VIC (BetaCoV/Australia/VIC01/2020, lineage B), IC19 (hCoV-19/England/IC19/2020, lineage B.1.13) and Alpha (hCoV-19/England/204690005/2020, lineage B.1.1.7) have been described previously ^11^. SARS-CoV-2 Delta (lineage B.1.617.2) and Omicron (lineage B.1.1.529/BA.1) isolates were a kind gift from Wendy Barclay (Imperial College London, UK) ^14, 31^. Viruses were propagated by infecting Caco2 cells at MOI 0.01 TCID50 per cell, in DMEM culture medium supplemented with 1% FBS and 1% penicillin/streptomycin, at 37 °C. Virus was collected at 72 hpi and clarified by centrifugation at 2100xg for 15 min at 4 °C to remove any cellular debris. Virus stocks were aliquoted and stored at −80°C. Virus stocks were quantified by extracting RNA from 100 μl of supernatant with 1 μg carrier RNA using Qiagen RNeasy clean-up RNA protocol, before measuring viral E RNA copies per ml by RT-qPCR as described previously ^27^.

### Live virus infections

Caco2 (1×10^5^ cells/well) and HeLa-ACE2 (6.5×10^4^ cells/well) were seeded in 24-well plates one day before infection. Cells were infected with 1000 E RNA copies per cell in 200ul culture medium. After 2h incubation at 37°C, cells were carefully washed with PBS to remove excess virus and fresh culture medium was added. For inhibition assays, cells were pre-treated with inhibitors at the indicated concentrations for 2h prior to infections and maintained throughout the experiment. At the indicated time points, cells were collected for analysis.

For Calu3 cell infections, 2×10^5^ cells/well were seeded into 12-well plates and grown until confluent. Where indicated, cells were pre-treated with indicated concentrations of recombinant human IFNγ (Peprotech) for 8h before cells were infected with 1000 E RNA copies SARS-CoV-2 per cell in 400μl culture medium. The inoculum was thoroughly washed off with PBS after 2h and fresh culture medium added. At 36 hpi, cells were harvested for protein lysates and flow cytometry. Culture supernatants from infected cells were clarified by centrifugation at 2100xg for 15 min at 4°C and viral E RNA copies measured by RT-qPCR. To determine infectivity of these viral supernatants, Caco2 and HeLa-ACE2 cells were pre-treated with 5 μM Ruxolitinib (Bio-Techne) for 1h to inhibit JAK signaling (from carryover interferon) and then infected with 1000 E copies/cell of virus as described above and harvested at 24hpi for flow cytometry analysis. Ruxolitinib was maintained throughout.

### Pseudovirus production

Spike pseudoviruses (PVs) were made by co-transfection of spike, p8.91 and pCSLW plasmids as described previously ^39^. To determine GBP inhibition, PV plasmids were transfected together with GBP vector or empty vector (EV) control expressing only BFP reporter. Briefly, 5×10^4^ 293T cells were seeded onto 24-well plates for 24h and then transfected with 260 ng p8.91, 260 ng pCSLW, 40 ng spike and 20-120 ng GBP vectors or EV control using Fugene6 (Promega). For larger scale production cells were seeded in 6-well plates with cell numbers and transfection reagents scaled up five-fold. For HIV-1 Env PV, 293T cells were seeded as described above and transfected with 240 ng p8.91, 240 ng pCSLW,120 ng pSVIII-JRFL Env and indicated doses of GBP or empty vector control.

PV supernatants were collected at 48 and 72 h post-transfection and purified through 0.45 μm centrifuge tube filters (Corning) or 0.45 μm syringe-filters (Starlab) and used within 24h without freeze-thawing. The amount of PV in the supernatant was determined by measuring the supernatant RT activity using SYBR-green–based product enhanced reverse transcription assay (SG-PERT) by qPCR, performed as described previously ^52^.

### Pseudovirus infection

Target cells were seeded into white 96-well plates 24 h before infection (Caco2, Vero.E6 and Vero.E6-TMPRSS2 cells seeded at 1.5×10^4^ cells/well, 293T-ACE2, HeLa-ACE2 and HeLa-TZMbl cells seeded at 1×10^4^ cells/well). Cells were infected equal doses (25 μl) of PV supernatant and incubated at 37°C for 48 h without changing the media. Luciferase expression (RLU) was measured at 48 h post-infection using BrightGlo substrate (Promega) according to manufacturer’s instruction on Glomax luminometer (Promega). For inhibitor studies cells were pre-treated before infections for 2 h with Camostat mesylate (Apexbio, 0.2-100 μM) or E64d (Focus Biomolecules, 0.2-25 μM). To obtain infectivity (RLU/RT) values, RLU values were normalized to supernatant RT activity (measured by SG-PERT assay).

### HIV-1 infection

293T cells were seeded in 24-well plates as described above and transfected with 120ng pNL4.3 and indicated doses of GBP or empty vector control using Fugene6 to produce infectious HIV-1 virus in presence or absence of GBPs. Virus-containing supernatant was harvested and RT activity measured as described above. HeLa-TZMbl were seeded into white 96-well plates 24 h before infection (1×10^4^ cells/well) and infected with 10 μl of virus supernatant/well. Luciferase activity was measured 48 h post-infection as described above.

### GBP inhibition assay

Spike PV, HIV-1 Env PV or HIV-1 virus were made in presence of increasing doses of GBP2/5 or their mutants or empty vector control as described above. Supernatant RT activity (RT units) was measured by SG-PERT assay to determine PV or virus content in supernatants. As indicated, different cell lines were infected with equal doses of PV/virus supernatants for 48 h and luciferase expression (RLU) was measured as describe above. Infectivity of the supernatant was determined as a ratio of RLU to RT units (RLU/RT) and normalized to empty vector control (no GBP), set at 100% for each PV or virus.

### IFITM inhibition assay

Caco2 cells, WT or stably expressing IFITM1/2/3 (described above), were infected with indicated spike PVs and luciferase expression was measured 48 h post-infection as described above. Infectivity (RLU/RT) was normalised to Caco2 WT control (100%).

Alternatively, spike PV were made in presence of GBP5 or EV control and used to infect Caco2 WT and IFITM expressing cells. Infection of Caco2-IFITM1/2/3 cells with live SARS-CoV-2 virus was done as described above.

### Immunoblotting

PV and live viruses were concentrated and purified from the supernatant by centrifugation (2h 13,000g, 4^0^C) through a 20% sucrose cushion and resuspended in 20 μl Leammli buffer. Cells were harvested, lysed in RIPA buffer and protein concentration determined by the BCA assay (Thermo Fisher Scientific). Twenty micrograms of cell lysate and an equal volume of purified virus were separated by SDS-PAGE, transferred to nitrocellulose membrane and membranes probed using the following primary antibodies: anti-SARS-CoV Spike (100% overlap with the SARS-CoV-2 epitope) (Invitrogen, PA1-41165, 1:1000), anti SARS-CoV nucleocapsid (N) protein monoclonal antibody (clone CR3009, gift from Laura McCoy, 0.5 μg/ml), anti-HA-tag (Biolegend, 16B12, 1:2000), anti-Gag p24 (CFAR, ARP432, 1:2000), anti-GBP5 (Cell Signaling, D3A5O, 1:1000), anti-GBP2 (Santa Cruz, G-9, 1:200), and anti-tubulin (Sigma, DM1A, 1:1000). Primary antibodies were incubated overnight at 4°C with agitation and detected with fluorescent secondary antibodies: anti-Rabbit IgG (IRDye 800CW, Abcam, ab216773, 1:10,000), anti-Mouse IgG (IRDye 680RD, Abcam, ab216776, 1:10,000) and anti-Human IgG (IRDye 800CW, Licor, 925-68078, 1:10,000), and imaged with an Odyssey Cxl Infrared Imager (Licor). Immunoblots were analysed with Image Studio Lite software.

### Flow cytometry

Cells were harvested and detached with Trypsin-EDTA (Thermo Fisher Scientific), stained with Zombie NIR live-dead dye (Biolegend) in PBS (1:500, 5min), washed and fixed with 4% formaldehyde (Sigma) for 20-30 min at RT. Cells infected with live SARS-CoV-2 virus were fixed with 4% formaldehyde for 1h at RT. Cells were then washed and permeabilised with Perm buffer (Biolegend) for 5 min at RT and stained with primary antibodies (20 min at RT): anti-HA-tag-APC (Biolegend, 16B12, 1:150), anti-Gag-PE (Beckam Coulter, KC57, 1:250). SARS-CoV-2 infection was detected by intracellular staining for nucleocapsid (N) protein using anti-SARS-CoV N monoclonal antibody (human, clone CR3009, gift from Laura McCoy, 1 μg/ml) and secondary anti-Human IgG-AF488 (Jackson labs, 1:400). For analysis of cell surface spike infection, 293T cells were harvested with 5 mM EDTA (Sigma), washed, stained with Zombie NIR and anti-SARS-CoV-2 Spike antibody from human convalescent sera (NIBSC, 20/130 1:200) on ice for 30 min. Primary antibodies were detected with the secondary anti-Human IgG-AF488 before fixation and intracellular stain as described above.

### Immunofluorescence Microscopy

Caco2 cells were fixed in 4%PFA (Sigma) for 15 minutes, followed by permeabilisation in 0.25%Triton-TX100 (Sigma) for 15 minutes. A blocking step was carried out for 1h at room temperature with 1%BSA (Sigma) and 0.1% Triton TX100 in PBS. Primary antibody incubation was carried out for 1h at RT with rabbit anti-HA antibody (H6908, Sigma-Aldrich) to label IFITM1, IFITM2 and IFITM3 and mouse anti-CD63 (IB5, a gift from M. Marsh, UCL) to label endogenous CD63. Primary antibodies were detected with secondary anti-rabbit AlexaFluor-488 and anti-mouse AlexaFluor-568 conjugates (Jackson Immuno Research) for 1h. All cells were labelled with Hoechst33342 (H3570, Thermo Fisher). Images were acquired using the WiScan® Hermes 7-Colour High-Content Imaging System (IDEA Bio-Medical, Rehovot, Israel) at magnification 40X/0.75NA. Three channel automated acquisition was carried out sequentially. Scale bars and RGB composite images were created in in FIJI ImageJ software package ^53^.

### Statistical Analysis

Statistical significance was calculated using Prism 9 (GraphPad Prism) using indicated statistical tests and significance was assumed when p < 0.05.

## Acknowledgements

This work was funded by Wellcome Investigator Award 108079 followed by 223065 to C.J.

G.J.T. was funded by Wellcome Senior Fellowship 108183 followed by Wellcome Investigator Award 220863. C.J. and G.J.T. were also funded by MRC/UKRI G2P-UK National Virology consortium (MR/W005611/1) and the UCL COVID-19 fund. We thank the G2P-UK National Virology Consortium; W. Barclay and T. Peacock at Imperial College London, UK; G. Screaton Oxford University, UK; Dalan Bailey at Pirbright Institute UK; Laura McCoy at University College London UK; J.E. Voss and D. Huang at Scripps Research Institute USA; Joe Grove at the Centre for Virus Research, Glasgow, UK; Katie Doores at King’s College London UK, and Daniel Sauter at Ulm University Medical Center, Ulm Germany for provision of reagents and variant isolates. We also acknowledge members of the G2P-UK Consortium, as well as the Jolly lab and Towers lab for helpful discussions.

## Author contributions

C.J. conceived the study. D.M., A-K.R., M.X.V.W., T.B., T.H., and L.T., designed and performed the experiments and analysed the data. D.M., A-K.R., and C.J. wrote the manuscript with contributions from all authors.

---

**Extended data Figure 1:**
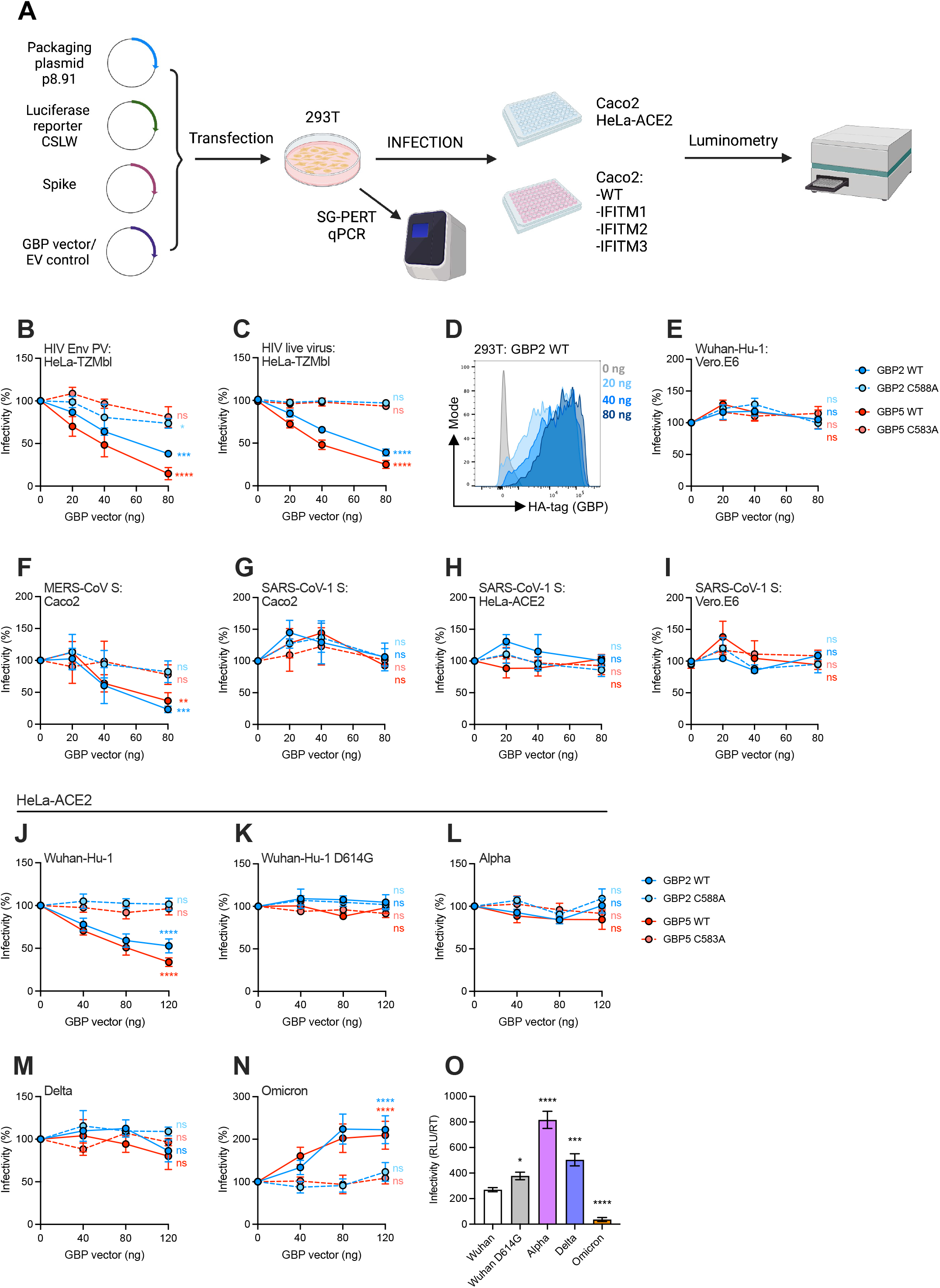
GBP restriction of HIV-1 Env, SARS-CoV-2, SARS-CoV-1 and MERS-CoV spike-mediated infection. **A)**Schematic of the pseudovirus (PV) infection assay. 293T cells were co-transfected with lentiviral packaging, luciferase reporter and spike plasmids (to make PV) and GBP vector or empty vector (EV) control. PV-containing supernatant was used to infect indicated cells lines and infection was quantified by measuring luciferase expression by luminometry. PV content in the supernatants was measured by SG-PERT qPCR assay. Created with BioRender.com B)HIV Env (JRFL) PV or **C)** HIV-1 live virus (NL4.3) were produced in 293T cells in presence of increasing amounts of plasmid encoding indicated GBPs or EV control. Infection was measured by luciferase assay (RLU) on HeLa-TZMbl reporter cells. Percentage infectivity of PV made in the presence of GBPs normalised to EV control (no GBP, set to 100%) are shown. **D)** Expression of HA-tagged GBP2 in PV-producing 293T cells measured by flow cytometry. Shown is a representative histogram, related to Fig. 1B. **E-I)** Spike PV were produced in presence of increasing doses of GBP2/5, GBP mutants or EV control and infection was measured on indicated cell lines (described in A). Percent infectivity was normalised to EV control for **E)** SARS-CoV-2 Wuhan-Hu-1 spike PV titrated on Vero.E6 cells, **F)** MERS-CoV spike PV titrated on Caco2 cells, and SARS-CoV-1 spike PV titrated on **G)** Caco2, **H)** HeLA-ACE2, and **I)** Vero.E6 cells. **J-O)** SARS-CoV-2 spike PV made in presence of increasing doses of indicated GBPs were titrated on HeLa-ACE2 cells (as described above). Shown is percent infectivity normalised to EV control for **J)** Wuhan-Hu-1, **K)** Wuhan-Hu-1 D614G, **L)** Alpha, **M)** Delta, and **N)** Omicron spike PV infection. **O)** Comparison of particle infectivity (RLU/RT) of spike PVs on HeLa-ACE2 cells made in the absence of GBPs. Shown is mean ±SEM from three independent experiments. A-N) Two-way ANOVA with Dunnett’s post-test. Asterisks denote statistical significance for GBPs (120 ng) compared EV control. (O) RM one-way ANOVA with Dunnett’s post-test. ns, not significant; *p < 0.05; **p < 0.01; ***p < 0.001; ****p < 0.0001.

**Extended data Figure 2:**
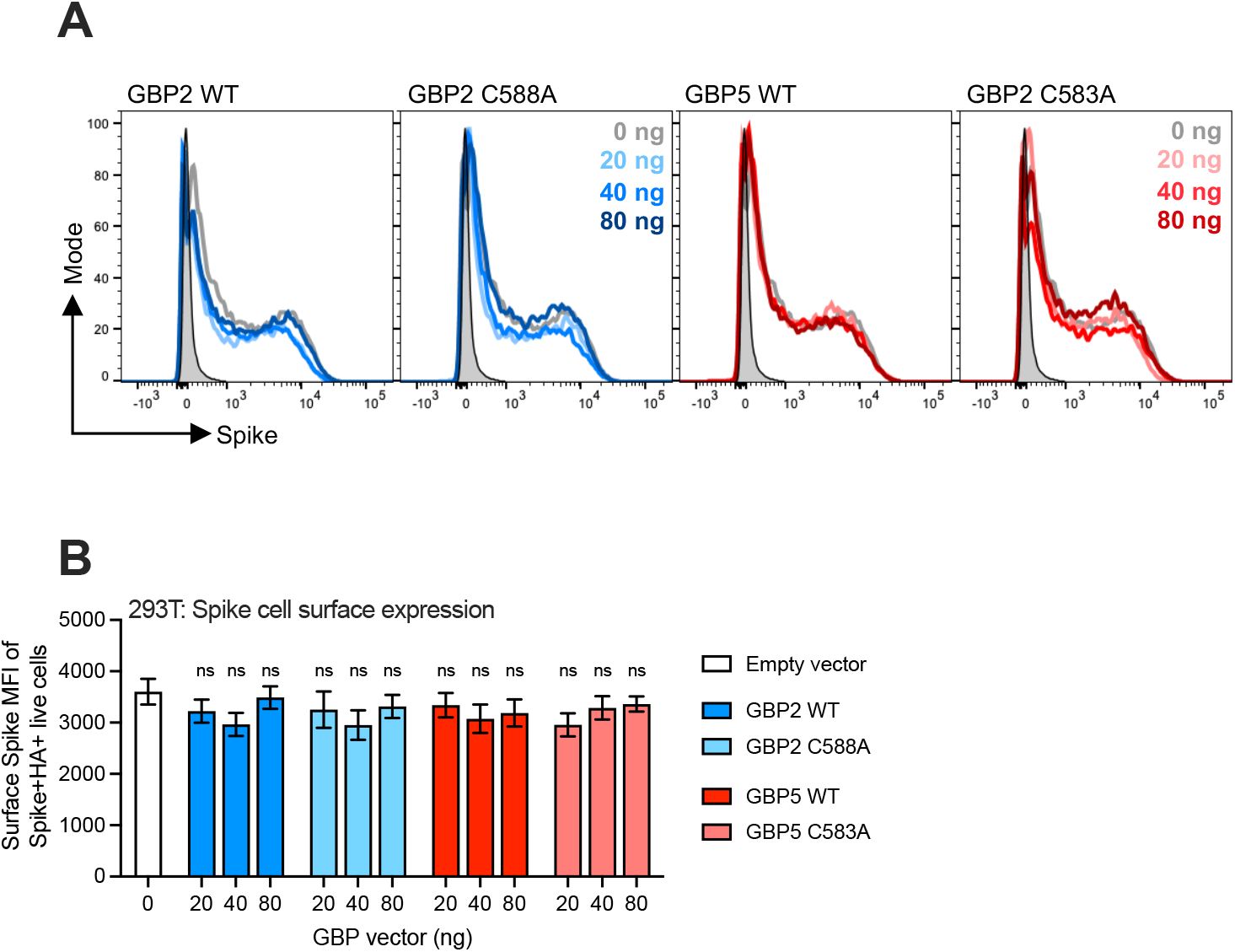
GBPs do not impact SARS-CoV-2 Spike cell surface expression. 293T cells were transfected with Wuhan-Hu-1 spike, lentiviral packaging and reporter plasmids and increasing doses of GBP2/5 or GBP mutants. Cells were stained for cell surface expression of spike and intracellular expression of HA-tagged GBPs. **A)** Shown are representative histograms of spike expression in HA+ live cell population. **B)** Cell surface expression of spike (MFI) in the Spike+HA+ live cell population. Shown is mean ±SEM from three independent experiments. Different GBP treatments were compared to EV control using a RM one-way ANOVA with Dunnett’s post-test. ns, not significant.

**Extended data Figure 3:**
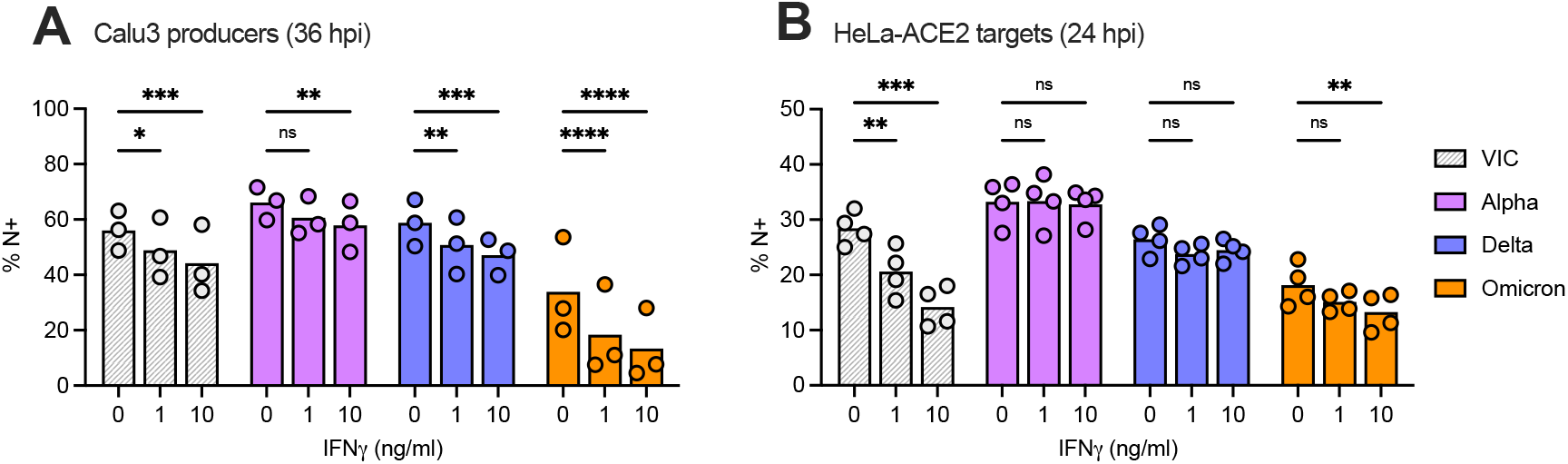
Effects of IFNγ treatment of Calu3 cells on virus infection and infectivity. **A)** Calu3 cells were pre-treated with indicated doses of IFNγ for 8 h and infected with indicated SARS-CoV-2 virus isolates for 36 h. Infection was measured by flow cytometry analysis to quantify N expression shown as % N+ cells. **B)** Equal doses of virus (measured by E copies) from the supernatant of IFNγ-treated Calu3 cells from A) were used to infect HeLa-ACE2 cells for 24 h. Infection was measured by flow cytometry. Mean and individual values from two independent experiments are shown. Two-way ANOVA with Dunnett’s post-test was used. ns, not significant; *p < 0.05; **p < 0.01; ***p < 0.001; ****p < 0.0001.

**Extended data Figure 4:**
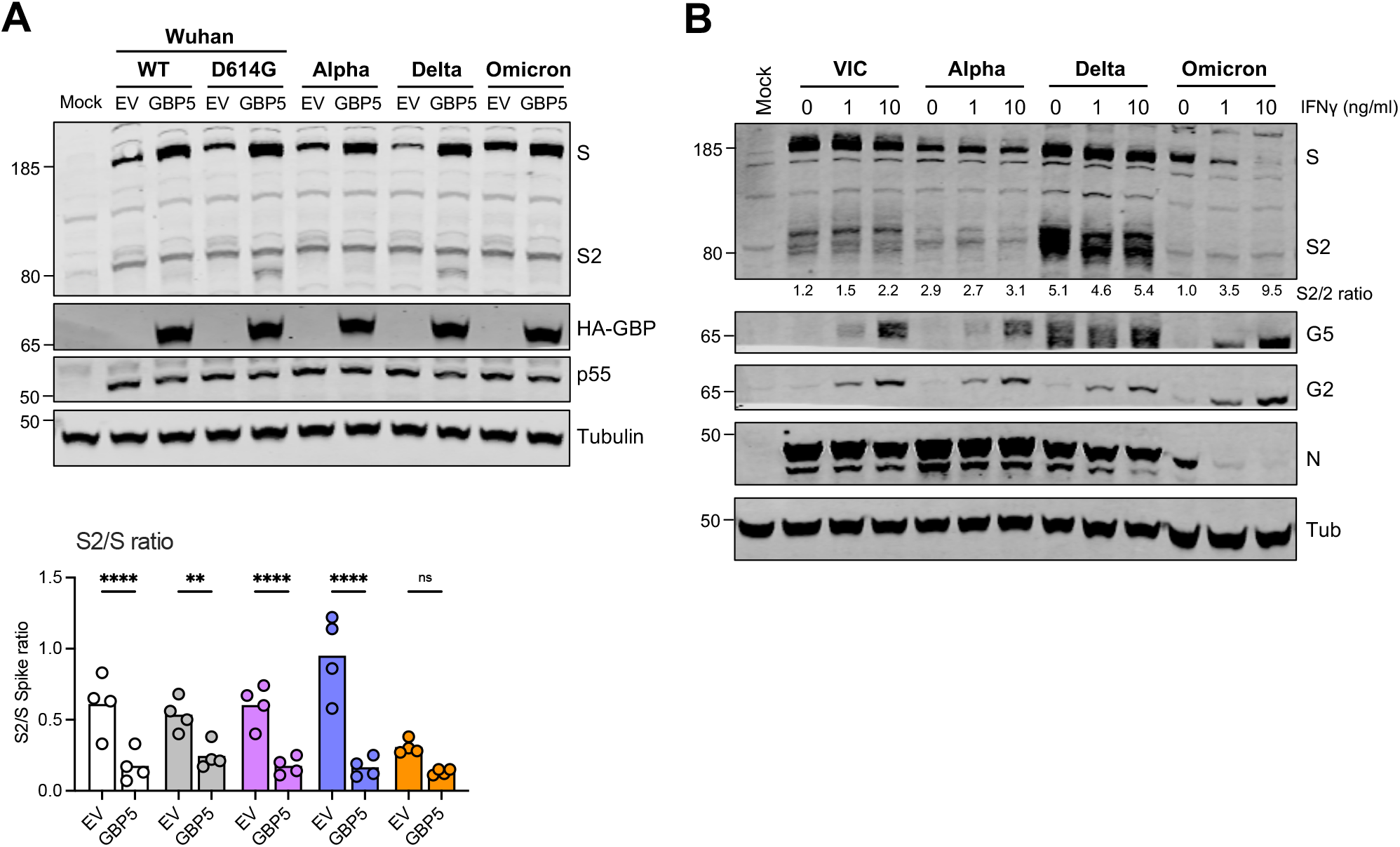
GBP2/5 and SARS-CoV-2 spike cleavage in cell lysates. **A)** VOC spike PVs were produced in the presence of 80 ng of GBP5 plasmid or EV control. Cell lysates were immunoblotted for spike, lentiviral Gag (p55), GBP5 (HA-tag) and tubulin. Shown is a representative immunoblot (corresponding PV immunoblot is shown in Fig. 2C). Graph shows quantification pooled from four independent experiments, measuring the relative proportion of cleaved spike (S2/S). Mean and individual values are shown. **B)** Calu3 cells were treated with indicated doses of IFNγ for 8 h and infected with indicated SARS-CoV-2 variants for 36 h. Cell lysates were immunoblotted for spike (S), GBP2, GBP5, nucleocapsid (N) protein and tubulin. Band quantification shows relative proportion of cleaved spike (S2/S). Corresponding virus immunoblot is shown in Fig. 2D. Two-way ANOVA with Dunnett’s post-test was used. ns, not significant; *p < 0.05; **p < 0.01; ***p < 0.001; ****p < 0.0001.

**Extended data Figure 5:**
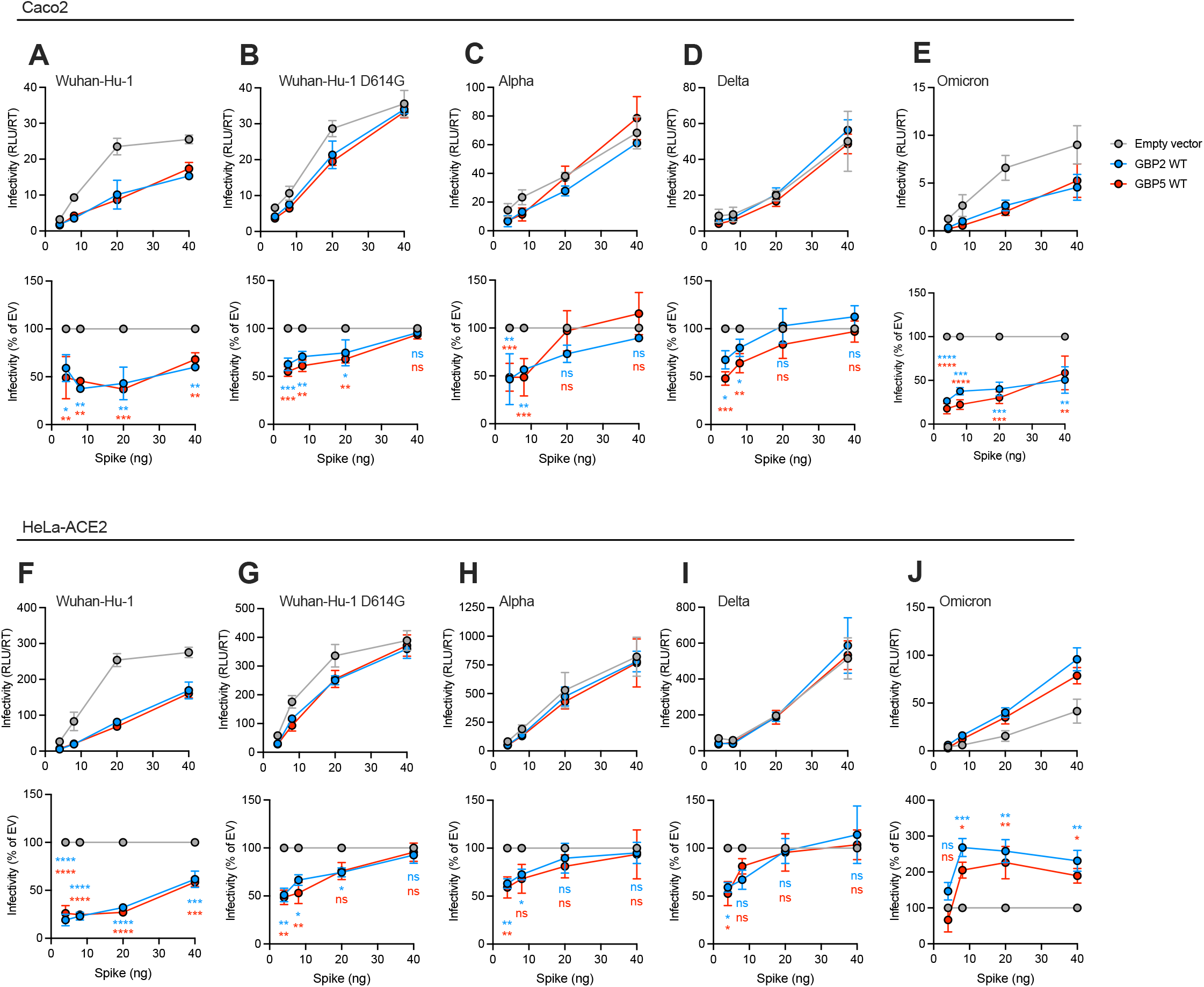
Reduced expression of spike restores GBP restriction. Spike PVs were produced with fixed amount of GBP2/5 vector or EV control (80 ng) and decreasing doses of spike plasmid. Upper panels show raw infectivity values (RLU/RT) and lower panels show percent infectivity of GBP2/5 PVs normalised to EV control at each dose of spike plasmid. **A)** Wuhan-Hu-1, **B)** Wuhan-Hu-1 D614G, **C)** Alpha, **D)** Delta, and **E)** Omicron spike PV titrated on Caco2 cells. **F)** Wuhan-Hu-1, **G)** Wuhan-Hu-1 D614G, **H)** Alpha, **I)** Delta, and **J)** Omicron spike PV titrated on HeLa-ACE2 cells. Shown is mean ±SEM from three independent experiments. Two-way ANOVA with Dunnett’s post-test was used. Asterisks denote statistical significance for GBP2/5 treatment compared EV control. ns, not significant; *p < 0.05; **p < 0.01; ***p < 0.001; ****p < 0.0001.

**Extended data Figure 6:**
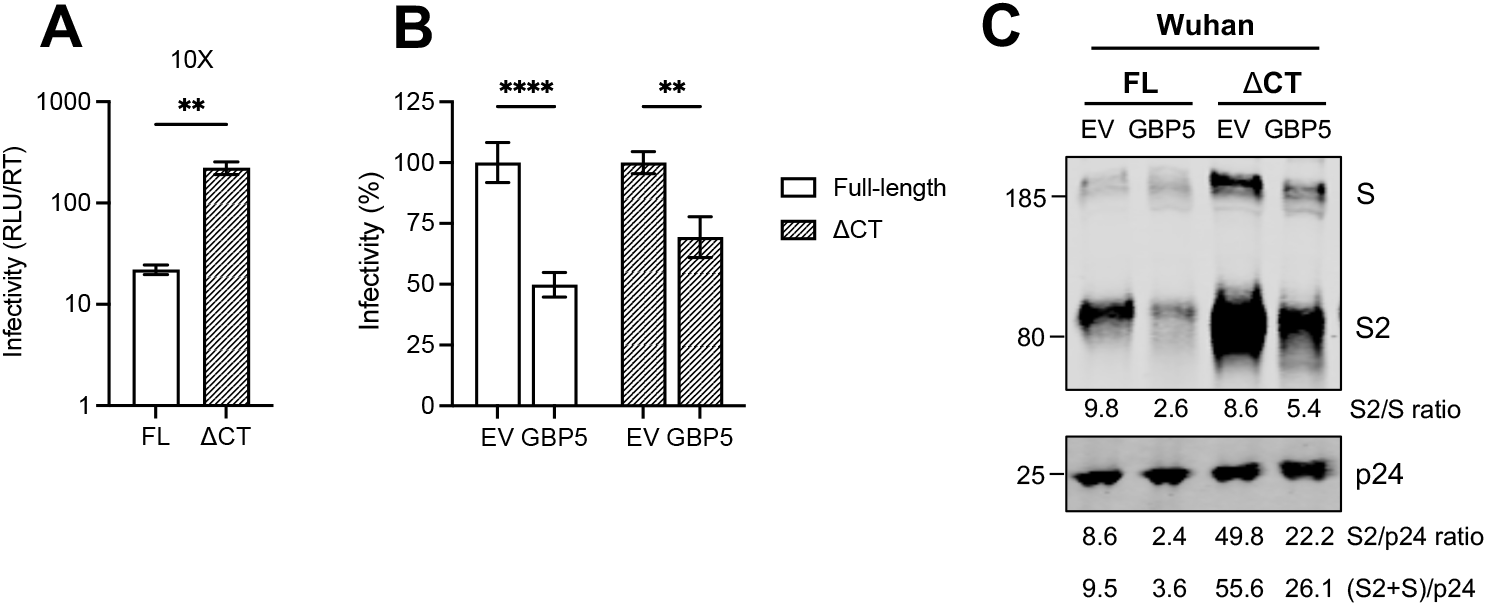
Increased Wuhan WT Spike incorporation into PV does not rescue from GBP restriction. Wuhan WT spike PV were produced in presence of GBP5 or EV control using plasmids encoding either full-length (FL) or a 19 residue C-terminal truncation (ΔCT) Wuhan spike sequence. Spike PV were titrated on Caco2 cells, showing **A)** raw infectivity (RLU/RT) values in absence of GBP5 expression (EV control) and **B)** percent infectivity normalised to EV control for each Wuhan spike (FL and ΔCT). **C)** Wuhan FL and ΔCT spike PVs produced the presence of GBP5 or EV control were purified from culture supernatants and immunoblotted for spike and lentiviral Gag (p24). Band quantification shows relative proportion of cleaved spike in PV (S2/S), cleaved spike incorporation into PV (S2/p24) and total spike incorporation ((S2+S)/p24).

**Extended data Figure 7:**
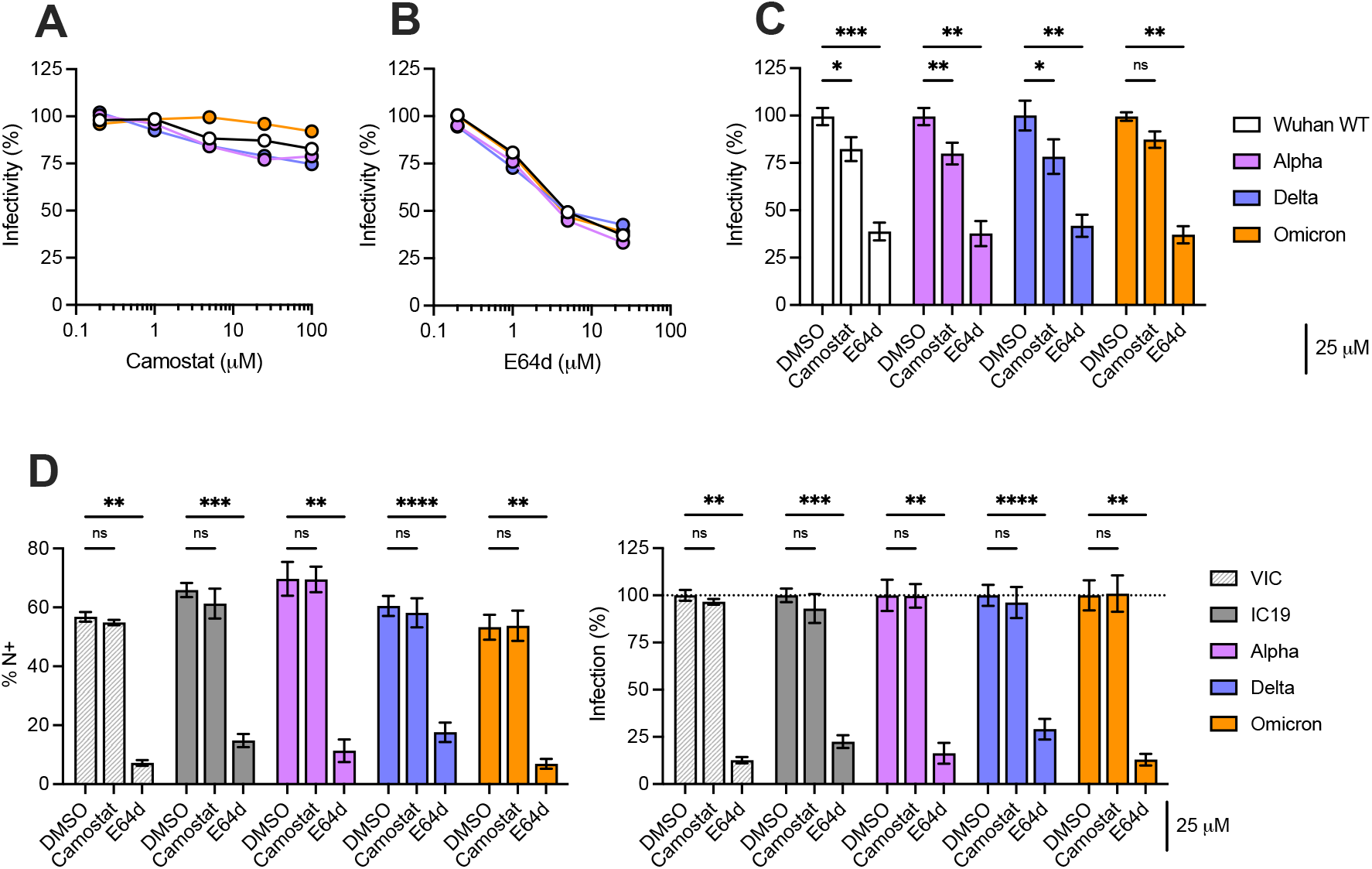
SARS-CoV-2 favors endosomal entry in HeLa-ACE2 cells. Inhibition of spike-mediated PV infection of HeLa-ACE2 cells by **A)** Camostat and **B)** E64d inhibitors. Shown is a representative titration. **C)** Inhibition of spike PV infection of HeLa-ACE2 cells in the presence of 25 μM Camostat or E64d normalised to DMSO control. Infection was measured by luciferase assay. **D)** Inhibition of SARS-CoV-2 live virus infection of HeLa-ACE2 cells in presence of 25 μM Camostat or E64d. Infection was measured by flow cytometry staining for nucleocapsid (N) protein and shown as the percentage N-positive cells (% N+). Left panel shows percentage N+ cells at 24 hpi, and the right panel shows the percentage infection for each virus normalized to the corresponding DMSO control. Shown is mean ±SEM from three independent experiments. Two-way ANOVA with Dunnett’s post-test was used. ns, not significant; *p < 0.05; **p < 0.01; ***p < 0.001; ****p < 0.0001.

**Extended data Figure 8:**
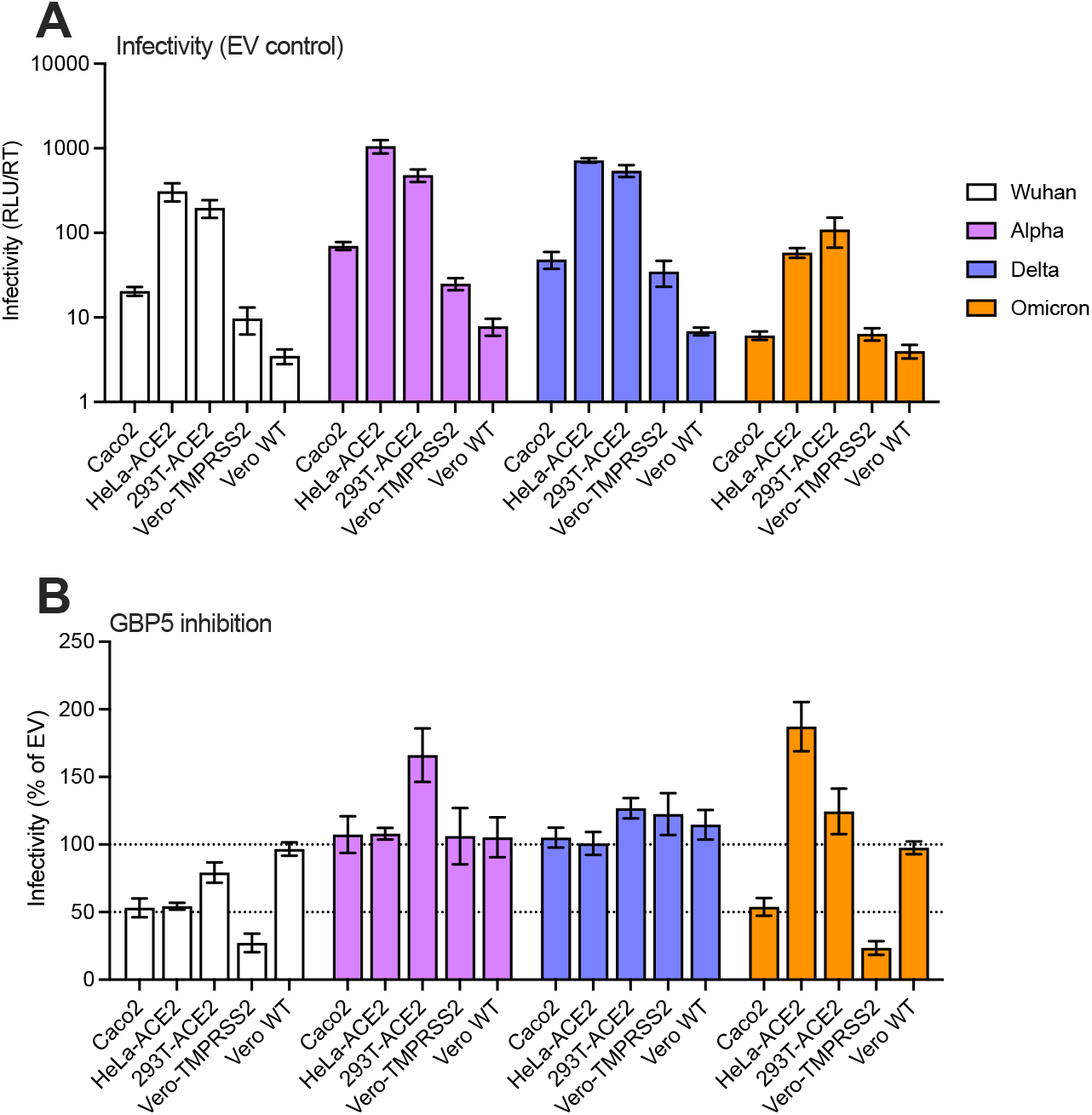
PV Infection and GBP5 inhibition across cell lines. Indicated spike PV were made in presence of 80 ng GBP5 or EV control and used to infect indicated cell lines: Caco2, HeLa-ACE2, 293T-ACE2, Vero.E6-TMPRSS2 and Vero.E6 WT. **A)** Shown are raw infectivity (RLU/RT) values for infection with PV made in absence of GBP5 (EV control). **B)** Shown is percent infectivity of PV made in presence of GBP5 normalised to EV control for each spike PV titrated on each cell line. Shown is mean ±SEM from three independent experiments.

**Extended data Figure 9:**
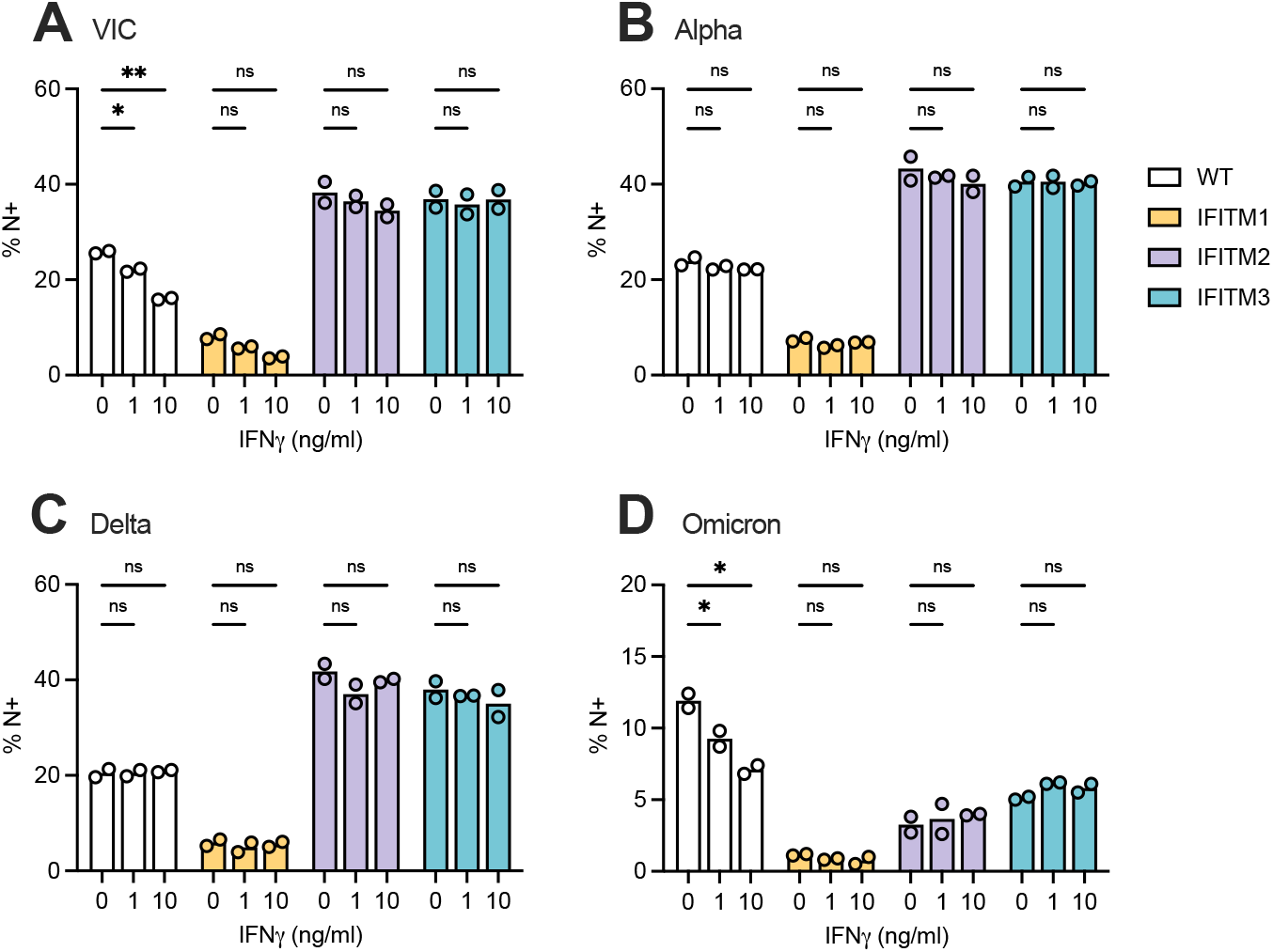
IFNγ treatment does not impact SARS-CoV-2 sensitivity to IFITMs. Calu3 cells were treated with indicated doses of IFNγ for 8 h to induce expression of GBP2/5 and infected with indicated SARS-CoV-2 variants. At 36 hpi, virus-containing supernatant was harvested. Equal doses of virus (measured by E copies) from the supernatants were used to infect WT or IFITM1/2/3 transduced Caco2 cells for 24 h. Infection was measured by flow cytometry staining for nucleocapsid (N) protein and shown as the percentage N-positive cells (% N+). Mean and individual values of two replicates from one experiment are shown. Two-way ANOVA with Dunnett’s post-test was used. ns, not significant; *p < 0.05; **p< 0.01; ***p < 0.001; ****p < 0.0001.

**Extended data Figure 10:**
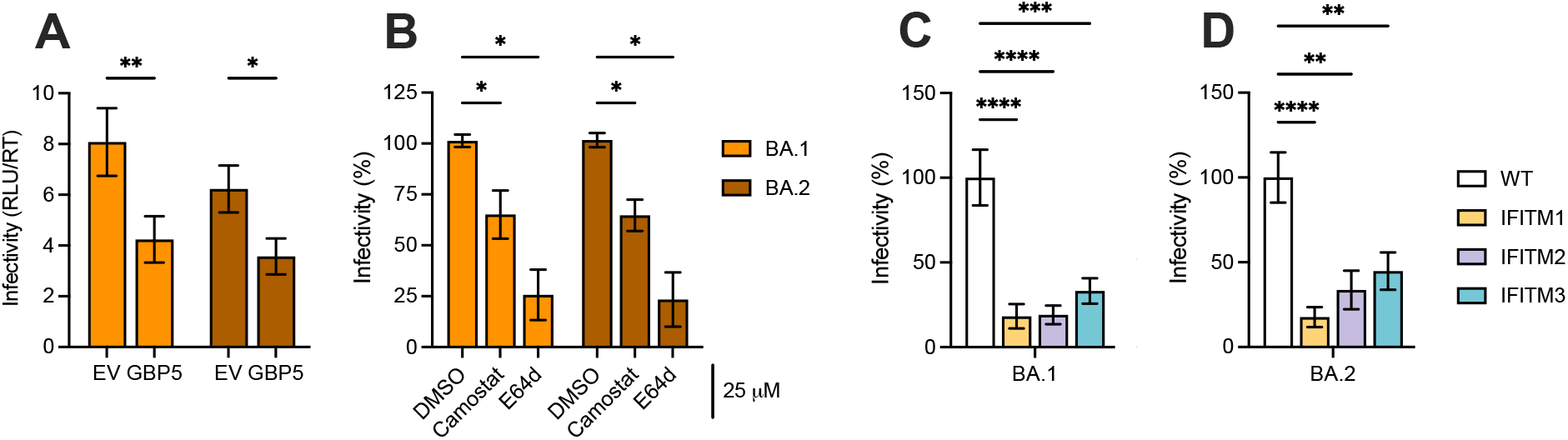
Omicron BA.1 and BA.2 spike PV are equally sensitive to GBP5, entry inhibitors and IFITMs. **A)** Omicron BA.1 and BA.2 spike PV were produced in presence of GBP5 or EV control and titrated on Caco2 cells. Shown are raw infectivity (RLU/RT) values. **B)** Inhibition of spike PV infection of Caco2 cells in the presence of 25 μM Camostat or E64d normalised to DMSO control. **C-D)** PV infection of IFITM transduced Caco2 cells with **C)** BA.1 and **D)** BA.2 spike PV. Data are shown as percent infectivity normalised to WT Caco2 cells (no IFITM over-expression). Bars show mean ±SEM from three independent experiments. Two-way ANOVA with Dunnett’s post-test (A, B) or one-way ANOVA (C, D) were used. ns, not significant; *p < 0.05; **p < 0.01; ***p < 0.001; ****p < 0.0001.

